# A structured study of cross-condition prediction of transcriptional responses to gene perturbations

**DOI:** 10.64898/2026.07.30.741892

**Authors:** Ouyang Zhu, Jun Li

## Abstract

Gene perturbation experiments coupled with transcriptomic profiling are crucial for uncovering causal gene-gene relationships, yet it remains cost-prohibitive to systematically explore perturbation responses across diverse biological conditions. As a result, in silico prediction of perturbation response has emerged as an important strategy for guiding cost-effective experimental design. Although recent methods have begun to address cross-condition perturbation prediction, it remains under-characterized across scenarios defined by whether the perturbation has been observed during training under other biological conditions. Here, we study cross-condition prediction under both seen- and unseen-perturbation scenarios. We introduce TranScouter, a lightweight encoderdecoder framework that represents perturbed genes using LLM-derived embeddings of their text summaries and represents biological conditions using transcriptomic profiles of control cells from the target condition. Across evaluated benchmarks, TranScouter performs competitively in both scenarios. We further use empirical analyses to characterize how condition-space coverage and perturbation-effect transferability shape cross-condition performance.

## 1 Introduction

Elucidating the structure and dynamics of gene-gene interactions is fundamental for understanding how genes interact to control cellular function and for developing targeted therapeutic strategies [1–4]. Genetic perturbation experiments have emerged as a powerful approach in this quest by systematically altering specific genes and observing the resulting changes in the expression levels of other genes [5–8]. Advances in CRISPR-Cas9, CRISPR interference and activation, and high-throughput single-cell screening platforms such as Perturb-seq have greatly expanded our ability to interrogate gene function and infer causal relationships at single-cell resolution [9–13].

Despite these technological advances, experimental exploration of gene perturbation effects remains fundamentally limited by scale. In addition to the large number of candidate genes that could be perturbed, the transcriptional outcome of a perturbation can differ across biological backgrounds. In this work, we use the term “biological condition” broadly to refer to the baseline context in which a perturbation is applied, including factors such as cell type, cell line, stimulation state, or treatment condition. Because the same perturbation may induce different transcriptional responses under different biological conditions, comprehensive characterization of gene function requires studying perturbations across many gene-condition pairs. This creates a vast combinatorial space of genes × conditions, making exhaustive experimental measurement both time-consuming and prohibitively expensive [14–16]. Computational models that can predict perturbation responses in silico therefore offer an important opportunity to prioritize experiments and guide downstream validation.

A growing number of computational methods have been proposed to predict transcriptional responses to perturbations [17–21]. A large body of this work has focused on prediction within a fixed biological condition, where the model generalizes across perturbation identities while the cellular context remains unchanged (Figure 1a). In this setting, methods such as GEARS, biolord, Scouter, and GenePert differ in multiple respects, including how they represent the perturbed gene, which draws on sources such as gene regulatory graphs, Gene Ontology structure, or text-derived gene embeddings. These studies establish the importance of transferable perturbation representations for perturbation-response prediction.

**Figure 1:**
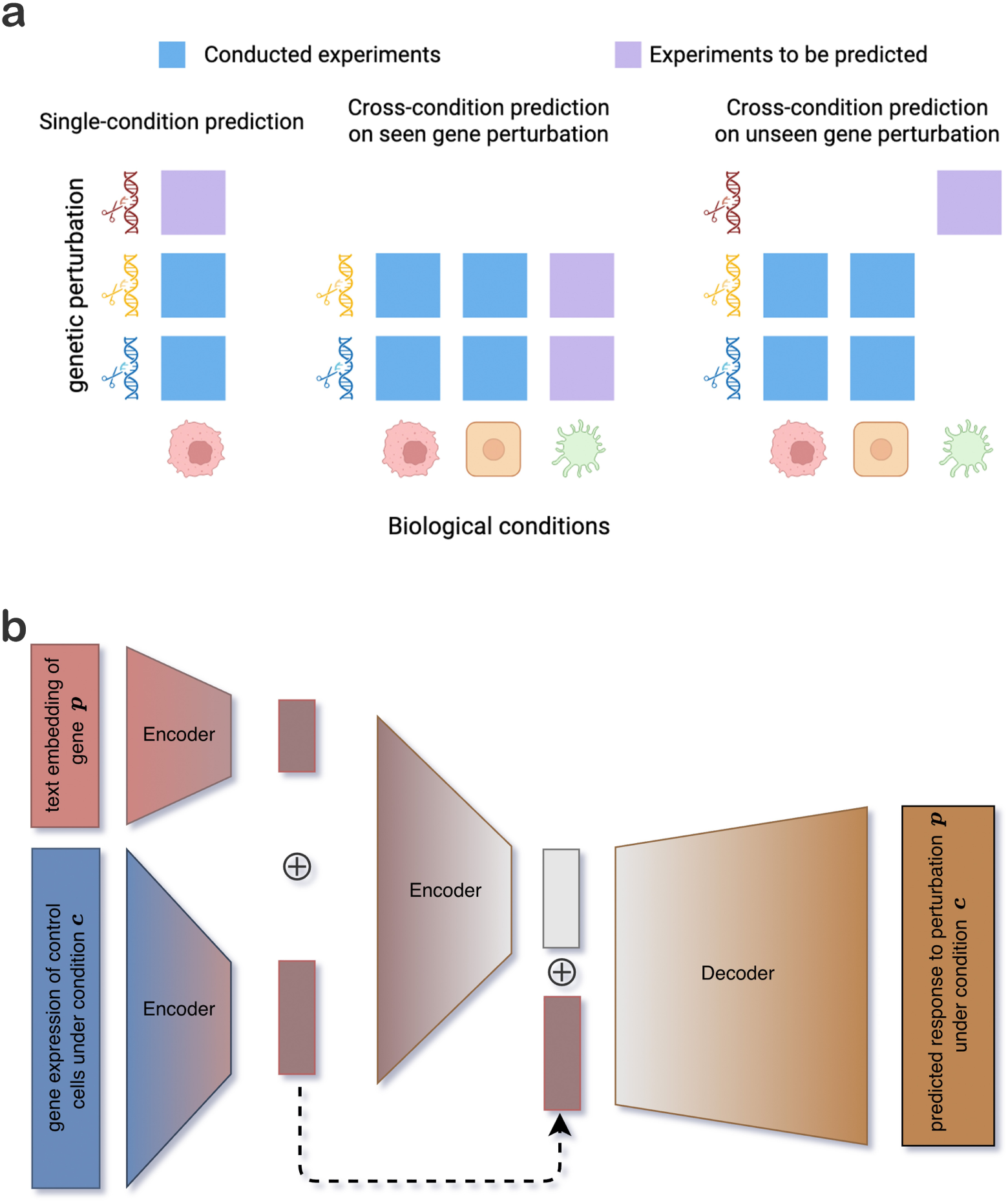
Overview of cross-condition perturbation prediction and the TranScouter framework. **a**, Schematic comparison of single-condition prediction, cross-condition prediction for seen perturbations, and cross-condition prediction for unseen perturbations. **b**, TranScouter architecture. A pretrained text-derived embedding of the perturbed gene and the control expression profile from the target biological condition are encoded separately, combined into a joint latent representation, and decoded to predict the post-perturbation expression profile in the target condition. The encoded condition representation is reintroduced near the bottleneck to preserve condition-specific information during decoding.

Beyond fixed-condition prediction, several methods have addressed cross-condition response prediction, including scGen [22], trVAE [23], TxPert [24], and STATE [25, 26]. These methods use different modeling strategies, but collectively show that predicting perturbation responses across biological conditions is an increasingly active direction. However, these studies also leave open an important distinction within cross-condition prediction: predicting perturbations that have been observed in other biological conditions differs from predicting perturbations whose effects have not been observed in any training condition. We refer to these as the seen- and unseen-perturbation scenarios, respectively. The latter requires extrapolation along both the perturbation and condition axes and has received less attention in prior cross-condition perturbation prediction studies. In addition, prior work has placed less emphasis on empirically characterizing factors associated with cross-condition performance, including condition-space coverage and perturbation-effect transferability.

In this work, we present a structured study of cross-condition prediction of transcriptional responses to gene perturbations under both seen- and unseen-perturbation scenarios. We view this task as a representation-based problem: prediction in held-out biological conditions requires both a representation of the perturbation target and a representation of the target condition. We introduce TranScouter, a lightweight encoder-decoder framework that represents perturbed genes using text-derived embeddings [19, 27–29] and represents biological conditions using transcriptomic profiles of control cells from the target condition [30, 31], allowing the two scenarios to be evaluated within a unified modeling framework. We benchmark TranScouter against existing methods adapted to each prediction scenario, together with empirical baselines, and further analyze factors associated with cross-condition performance, including condition-space coverage and perturbation-effect transferability. Together, this study provides a lightweight framework and empirical insights that may help guide future work on cross-condition perturbation prediction.

## 2 Results

### 2.1 Overview of the TranScouter model

A central requirement for cross-condition prediction is that the target biological condition can be represented in a way that allows meaningful comparison with conditions observed during training. This is analogous to the setting studied in fixed-condition prediction methods such as Scouter and GenePert, where generalization becomes possible when perturbed genes are represented by informative continuous embeddings. Following the same logic, TranScouter formulates cross-condition prediction as a dual-representation framework: perturbations require a transferable representation of perturbation identity, and biological conditions also require transferable representations. In the present study, we instantiate this framework by representing perturbed genes using LLM-derived embeddings of gene function text and by representing biological conditions using the transcriptomic profiles of control cells from those conditions. From a transcriptional perspective, this condition representation captures key aspects of the biological background, including cell identity and baseline state, and can therefore provide an anchor for extrapolating perturbation effects to new conditions.

Based on this idea, TranScouter predicts post-perturbation transcriptional profiles in heldout biological conditions under both seen- and unseen-perturbation scenarios. Given as input a perturbed gene and the control-cell transcriptomic profile from a target condition, TranScouter predicts the corresponding post-perturbation gene expression vector for that condition through a lightweight encoder-decoder architecture (Figure 1b). A perturbation encoder maps the fixed LLM embedding of the perturbed gene into a learned latent representation, while a condition encoder maps the control-cell expression profile into a representation of the target biological condition. These two representations are combined and passed through an additional encoder to form a joint latent state containing both perturbation-specific and condition-specific information. The encoded condition is then reintroduced near the bottleneck to help preserve target-condition information during decoding [32]. Finally, a decoder outputs a predicted expression profile, reflecting the anticipated post-perturbation response given the input gene perturbation applied to the target biological condition. The model is trained end-to-end with a direction-aware loss [17] that encourages accurate prediction of both the magnitude and direction of gene expression changes.

### 2.2 Model Performance

We evaluated TranScouter on Jiang24 [33], a large-scale CRISPRi Perturb-seq dataset containing 1.6 million cells across 30 biological conditions defined by 6 cell lines (A549, BxPC-3, HAP1, HT-29, K562, MCF-7) and 5 cytokine treatments (IFNB, IFNG, INS, TGFB, TNFA). Each condition contains a condition-specific subset of gene perturbations, yielding 218 unique target genes and 1,656 observed perturbation-condition pairs after preprocessing. This multicondition design provides a useful benchmark for evaluating whether perturbation-response models can generalize to held-out biological conditions, with condition structure and perturbation coverage summarized in Supplementary Note 1.

To evaluate held-out-condition prediction, we split the data at the level of biological conditions. In each split, a subset of conditions was withheld from model training and used for evaluation, while control profiles from the held-out conditions were provided at test time as representations of the target biological state. Test cases were separated into the two scenarios defined above: a seen-perturbation scenario, in which the perturbation had been observed in at least one training condition, and an unseen-perturbation scenario, in which the perturbation was absent from all training perturbation-condition pairs. This design allowed us to evaluate cross-condition transfer when perturbation-specific response information was available elsewhere, as well as extrapolation when such information was unavailable.

Predictions were compared with ground-truth post-perturbation expression profiles using two primary metrics computed over differentially expressed genes (DEGs): mean squared error (MSE), which measures expression-magnitude error, and directional mismatch rate (DMR), which quantifies the fraction of DEGs whose predicted direction of change is opposite to the observed direction. We focus on MSE and DMR because they directly assess the magnitude and sign of DEG-level responses, which are central to biological interpretation of perturbation effects and have been widely used in perturbation-response modeling studies [17–19, 22]. Pearson correlation coefficient (PCC) is reported in Supplementary Note 2 as a complementary diagnostic of prediction performance.

We first evaluated the seen-perturbation scenario, in which the target perturbation had been observed in at least one training condition but not in the held-out target condition. This scenario tests whether perturbation-response information observed in other biological conditions can be transferred to a new condition. We compared TranScouter with scGen, trVAE, and an averaging baseline. In this baseline, the prediction for perturbation *p* in target condition *c^∗^* is defined as the average observed profile of the same perturbation across training conditions:

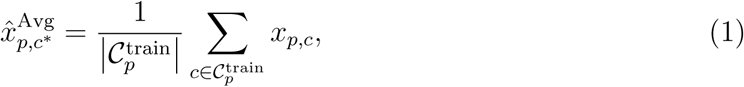

where *x_p,c_* is the average expression profile for perturbation *p* in condition *c*, and 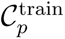 denotes the set of training conditions in which perturbation *p* is observed. This provides a simple reference that uses perturbation-specific information available in training but does not explicitly model the target biological condition.

Results are shown in Figure 2a, where each panel corresponds to one cell line and displays five cytokine treatments along the x-axis. The bars indicate the median MSE for each method, enabling side-by-side comparison across held-out condition settings. Overall, TranScouter achieved lower MSE than the evaluated methods across most conditions. Aggregated over all biological conditions, TranScouter achieved a median MSE of 0.15, compared to 0.20 for scGen, 0.20 for trVAE, and 0.68 for the averaging baseline, representing reductions of 22%, 23%, and 77%, respectively. In conditions such as HAP1 treated with IFNG or HT29 treated with IFNB, TranScouter showed larger margins over the evaluated methods, whereas the averaging baseline performed substantially worse, consistent with the limitation of averaging perturbed profiles without adapting to the target-condition baseline.

**Figure 2:**
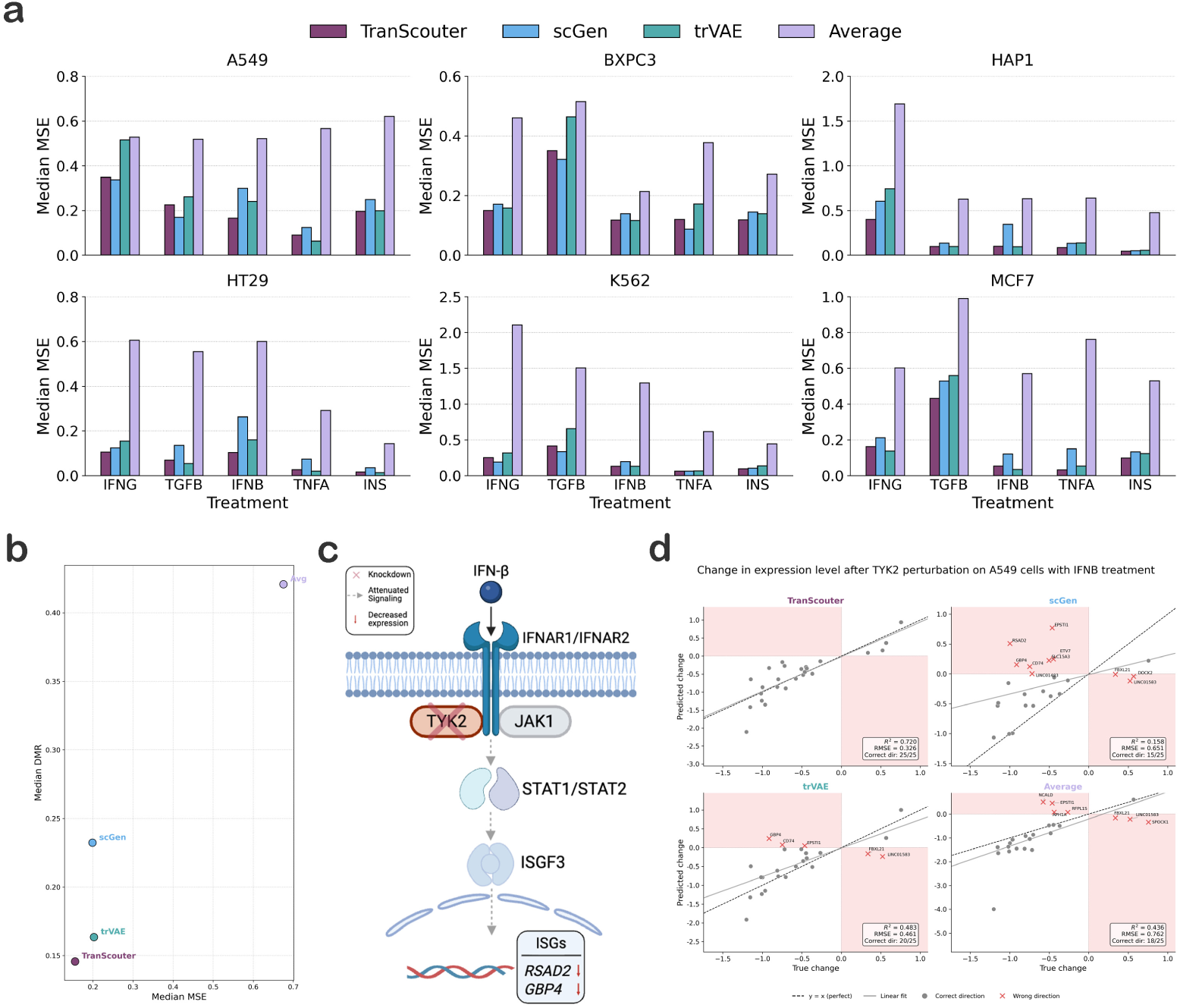
Cross-condition prediction performance in the seen-perturbation setting. **a**, Median mean squared error (MSE) over differentially expressed genes (DEGs) for TranScouter, scGen, trVAE, and the averaging baseline across 30 biological conditions. Each panel corresponds to one of six cell lines, with five cytokine treatments along the x-axis. **b**, Overall comparison of median MSE and directional mismatch rate (DMR) across held-out conditions. **c**, Biological schematic for TYK2 knockdown under IFN-*β* stimulation. **d**, Gene-level comparison of predicted versus observed expression changes for TYK2 knockdown in A549 cells under IFN-*β* treatment. Each point represents one DEG. Red shaded quadrants indicate predictions with the opposite sign from the true expression change. Panel annotations report squared Pearson correlation *R*^2^, root mean square error (RMSE), and the number of DEGs with correctly predicted direction.

In addition to MSE, we evaluated each method using DMR, with results summarized in Figure 2b. TranScouter achieved the lowest median DMR, reaching 0.14 across all conditions, compared with 0.23 for scGen, 0.16 for trVAE, and 0.42 for the averaging reference. trVAE showed reasonably strong directional performance in many conditions but was substantially more computationally expensive in our implementation: whereas scGen and TranScouter completed training in under 25 minutes on CPU, trVAE required more than 76 hours on average, and its GPU implementation did not finish within the 7-day runtime (see Methods for details).

To provide a concrete biological illustration, we examined the perturbation of TYK2 under IFN-*β* stimulation in A549 cells, a condition that strongly activates type I interferon signaling. TYK2 is a Janus kinase required for efficient signal transduction downstream of the type I interferon receptor, as illustrated in Figure 2c, and its knockdown is therefore expected to attenuate induction of many interferon-stimulated genes (ISGs) under IFN-*β* stimulation [34–36]. As shown in Figure 2d, TranScouter closely matched the true perturbation-induced expression shifts and correctly recovered the direction of change for all 25 plotted differentially expressed genes, whereas scGen and trVAE produced substantially more wrong-direction predictions, including several canonical ISGs. For example, RSAD2 (also known as viperin), a well-established antiviral effector induced robustly by type I interferons, was strongly down-regulated in the true data following TYK2 knockdown, reflecting the loss of interferon receptor signaling. TranScouter correctly predicted this down-regulation, but scGen instead predicted an up-regulation, which is inconsistent with the expected biology of TYK2 loss under IFN-*β* stimulation. Similarly, GBP4, a member of the guanylate-binding protein family involved in interferon-mediated host defense, was also down-regulated in the true data. TranScouter again captured the correct negative direction, whereas trVAE and scGen predicted the opposite sign, predicting a direction inconsistent with the expected attenuation of GBP4 induction following TYK2 loss. A similar pattern was observed for IFNAR2 perturbation, as demonstrated in Supplementary Note 3.

Taken together, the seen-perturbation results show that perturbation-response information observed in training conditions can support prediction in held-out biological conditions, and that combining perturbation representations with target-condition information provides an effective lightweight modeling strategy for this scenario.

We next evaluated the unseen-perturbation scenario. Unlike the seen-perturbation scenario, this setting does not provide observed response profiles for the target perturbation in other biological conditions, and therefore requires extrapolation along both the perturbation and condition axes. We compared TranScouter with three adapted reference approaches. First, we included CellOracle [37], a gene-regulatory-network-based method originally developed for simulating transcription factor perturbations and cell-state transitions. Because CellOracle relies on transcription-factor regulatory programs, it is applicable only to TF perturbations. In our dataset of 218 perturbations, 77 involve TF genes and are therefore within the scope of CellOracle, while the remaining perturbations are not directly available for CellOracle-based prediction. Second, we included an averaging baseline computed in the same way as in the seen-perturbation scenario. This baseline is not a valid unseen-perturbation predictor: by construction, the target perturbation is masked from all training conditions, whereas the baseline is computed from the masked same-perturbation profiles in other biological conditions. It therefore has access to perturbation-specific information deliberately withheld from TranScouter, and could not be computed for a perturbation never measured in any condition. We report it as an optimistic reference rather than as a deployable unseen-perturbation method. Finally, we implemented a correlation-based heuristic proposed in earlier work [38]. This method first constructs a gene–gene Pearson correlation matrix from the training data, where each entry reflects the Pearson correlation coefficient between two genes. Upon knockdown of a gene, its expression is set to zero, and the expression of correlated genes is adjusted in proportion to their pairwise correlations. Although correlation does not imply causal regulation, this heuristic provides a simple reference that can be applied without using observed responses of the held-out perturbation, in contrast to the optimistic averaging baseline.

Condition-specific MSE results are shown in Figure 3a, with one panel per cell line. Missing bars for CellOracle indicate cases where the held-out perturbations were not transcription factors and therefore fell outside the scope of CellOracle simulation. Across most conditions, TranScouter achieved the lowest error among methods evaluated on the full unseen-perturbation test set. For example, in BxPC-3 cells treated with IFNB, TranScouter obtained an MSE of 0.02 compared with 0.44 for the correlation-based heuristic and 0.26 for the optimistic averaging baseline; CellOracle, where applicable, obtained an MSE of 0.65 on the TF-only subset. In more difficult settings such as MCF-7 treated with TGFB, all methods incurred higher error, but TranScouter remained competitive at 0.53 compared with 1.19 for the correlation-based heuristic and 1.24 for the optimistic averaging baseline. When aggregated across all conditions, TranScouter achieved an average MSE of 0.24, compared with 0.55 for the correlation-based heuristic and 0.76 for the optimistic averaging baseline. CellOracle achieved an average MSE of 0.27, but this value was computed only over test perturbations involving TF genes, providing a narrower evaluation than the full unseen-perturbation benchmark. DMR showed a similar pattern, as summarized in Figure 3b. TranScouter achieved an average DMR of 0.18 across the full unseen-perturbation test set, compared with 0.44 for the correlation-based heuristic and 0.41 for the optimistic averaging baseline. CellOracle achieved a DMR of 0.36 on the TF-only subset where it was applicable. These results indicate that the representation-based strategy used by TranScouter can support cross-condition prediction even when no observed response profiles are available for the target perturbation during training.

**Figure 3:**
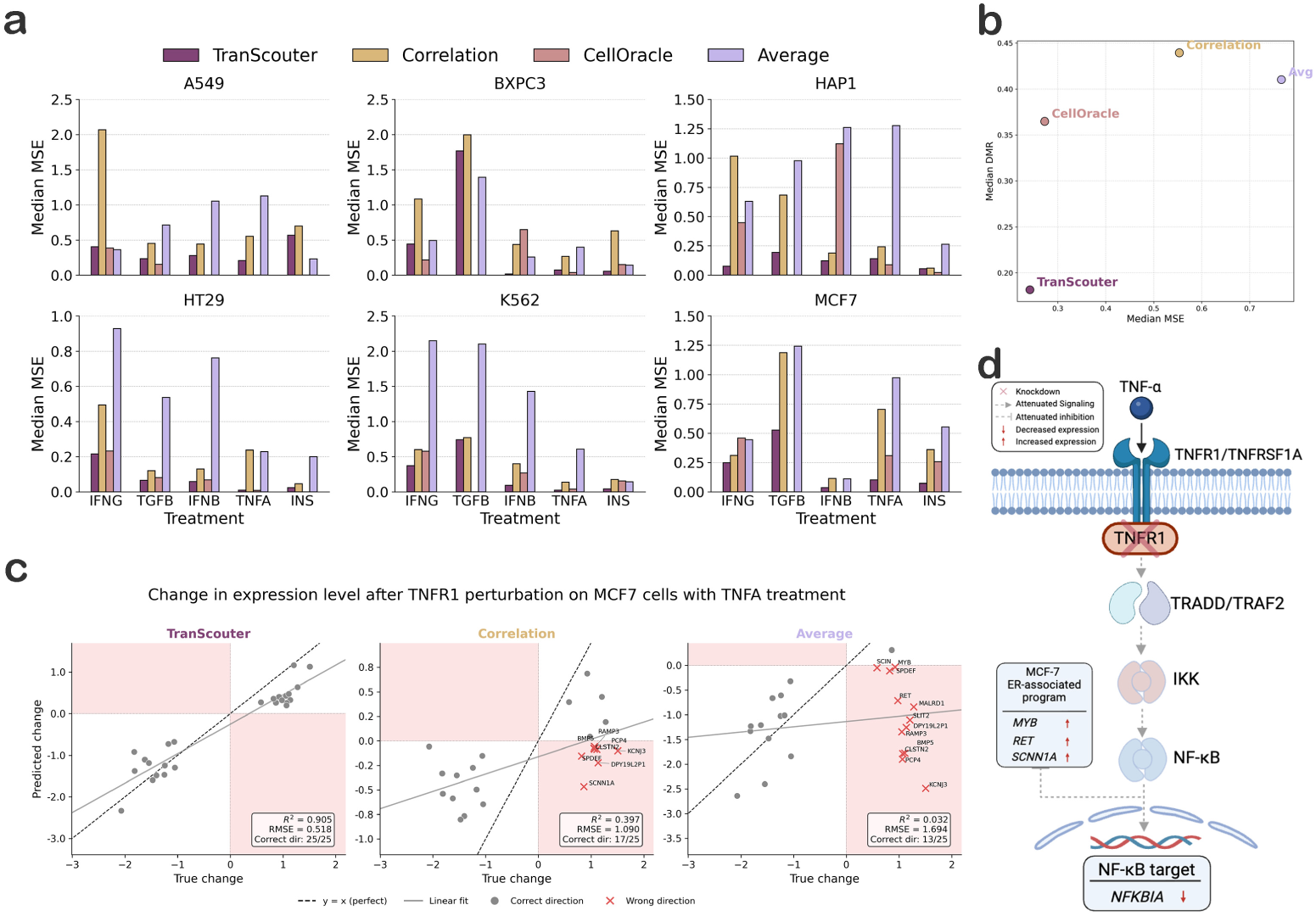
Cross-condition prediction performance in the unseen-perturbation setting. **a**, Median MSE on DEGs for each held-out biological condition in Jiang24, grouped by cell line and cytokine treatment. TranScouter is compared with a correlation-based heuristic, CellOracle, and a perturbation-averaging reference. CellOracle is evaluated only where the held-out perturbation target is within its TF/regulatory-gene scope; missing bars indicate conditions where CellOracle predictions were not available. **b**, Overall comparison of median MSE and DMR across held-out conditions. TranScouter is evaluated on the full unseen-perturbation benchmark, whereas CellOracle is evaluated on the subset of applicable perturbations. **c**, Predicted versus observed expression changes for TNFR1 perturbation in MCF-7 cells under TNF-*α* treatment. Each point represents one DEG. Red shaded quadrants indicate predictions with the opposite sign from the true expression change. Panel annotations report squared Pearson correlation *R*^2^, root mean square error (RMSE), and the number of DEGs with correctly predicted direction. **d**, Biological schematic for TNFR1 knockdown under TNF-*α* stimulation in MCF-7 cells.

To further examine model performance in the unseen-perturbation scenario, we examined TNFR1 knockdown in MCF-7 cells under TNF-*α* stimulation, as shown in Figure 3c. TNF-*α* normally signals through TNFR1 to activate the NF-*κ*B pathway, which in turn induces a set of immediate-early and feedback genes that regulate inflammatory responses. Among the most canonical targets is NFKBIA, encoding I*κ*B*α* protein, a negative feedback regulator that restrains NF-*κ*B activity. When TNFR1 is knocked down, this pathway is attenuated, and induction of NFKBIA is correspondingly reduced, as illustrated in Figure 3d. Consistent with this mechanism, the ground truth shows NFKBIA as strongly down-regulated, and while all methods captured the direction correctly, TranScouter produced predictions closest in magnitude to the truth, demonstrating higher fidelity to the expected treatment effect. Beyond such treatment-related genes, there are additional DEGs whose regulation reflects the joint influence of the treatment and the cell line. MCF-7 is an estrogen receptor (ER)-positive breast cancer cell line, in which the ER pathway governs the expression of many luminal-associated genes such as MYB, RET, SCNN1A, RAMP3, SPDEF, and KCNJ3. Prior studies have shown that TNF-*α* signaling through NF-*κ*B can antagonize ER activity in MCF-7 cells [39–42]. Accordingly, when TNFR1 is knocked down and TNF signaling is attenuated, ER-driven transcriptional programs may be de-repressed. This is consistent with the ground truth: ER-related genes such as MYB, RET, and SCNN1A are up-regulated, and TranScouter correctly predicts the positive direction of change, whereas other methods frequently make the wrong directional prediction. This example illustrates the type of structure that must be extrapolated in the unseen-perturbation scenario: the predicted response depends not only on the perturbed gene, but also on the cytokine treatment and cell-line background of the held-out condition. A similar qualitative pattern was observed for IFNGR2 knockdown in HT-29 cells under IFN-*γ* treatment (Supplementary Note 3), where interferon-stimulated genes fall as expected while cell-line-associated features shift in a manner consistent with the measured response.

### 2.3 Analysis of cross-condition generalization

Having evaluated both seen- and unseen-perturbation scenarios, we next examined factors associated with variation in cross-condition performance. We considered three complementary sources: condition-space coverage, including both training-condition diversity and target-condition proximity; perturbation-effect transferability across conditions; and modeling choices required to align perturbation responses with their condition-specific baselines.

#### Effect of training condition diversity

We first examined how performance depends on the number of distinct biological conditions included during training, using the unseen-perturbation scenario and reporting prediction error with MSE unless otherwise specified. TranScouter was trained on subsets of 5, 10, 15, 20, or 25 conditions and evaluated on heldout conditions. As shown in Figure 4a, with at least ten training conditions, performance was comparatively stable, with median MSE ranging from 0.12 to 0.16 across 10, 15, and 20 conditions, and 0.10 at 25 conditions. When training diversity collapsed to only five conditions, error rose sharply to 0.22. This pattern suggests that cross-condition prediction benefits from a minimally diverse set of training conditions, after which performance becomes comparatively stable; when too few conditions are available, it becomes harder for the model to separate perturbation-specific effects from baseline condition state.

**Figure 4:**
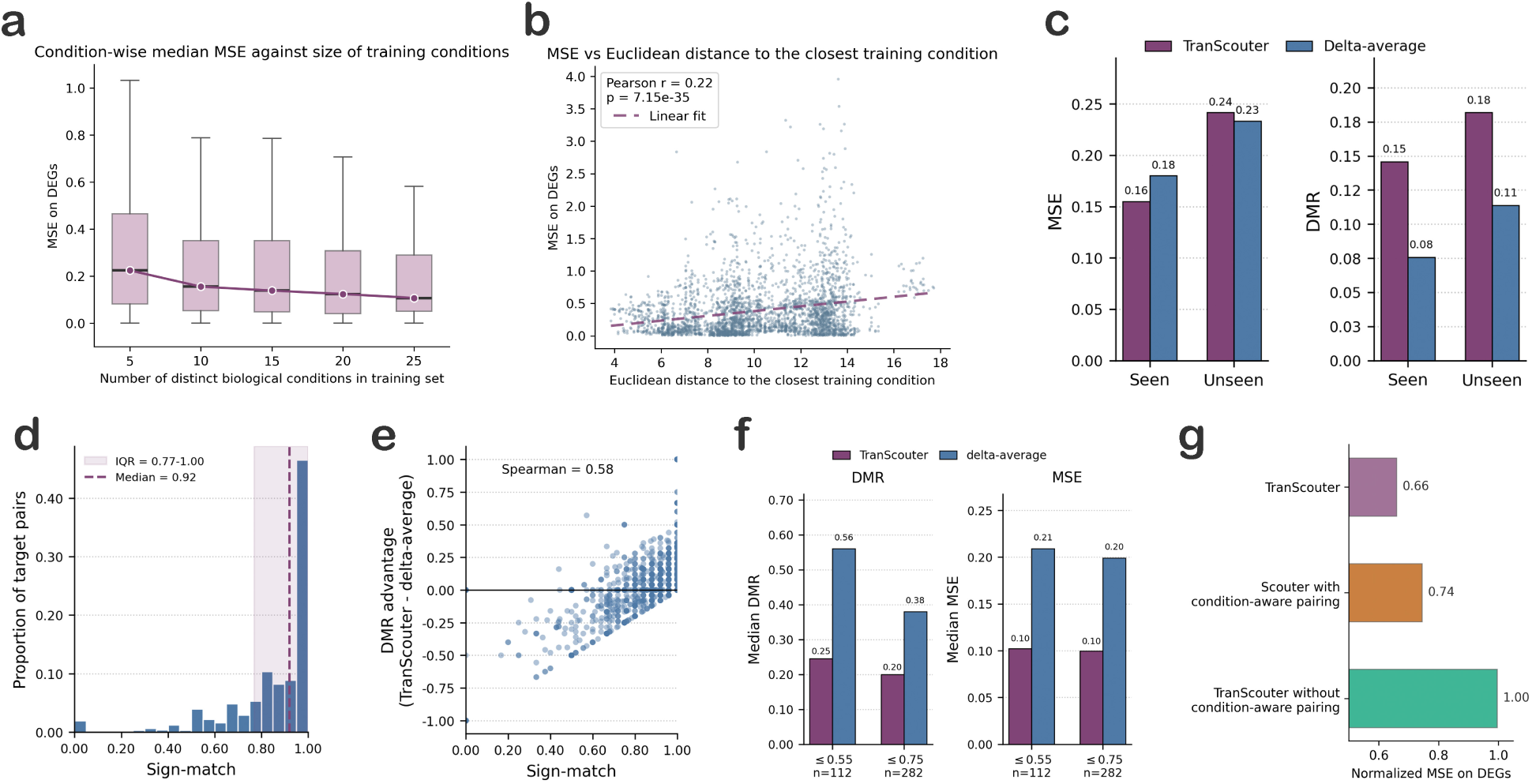
Analyses of factors associated with cross-condition generalization. **a**, Effect of training condition diversity on prediction accuracy. TranScouter was trained with 5, 10, 15, 20, or 25 biological conditions and evaluated on held-out conditions. **b**, Relationship between prediction error and distance from the held-out condition to the nearest training condition in the learned condition-embedding space. Each point represents a held-out perturbationcondition pair. **c**, Overall MSE and DMR comparison between TranScouter and the deltaaverage diagnostic in the seen- and unseen-perturbation scenarios. **d**, Distribution of signmatch scores across Jiang24 target perturbation-condition pairs. **e**, Relationship between sign-match and the DMR advantage of delta-average relative to TranScouter, defined as DMR_TranScouter_ − DMR_Δavg_. **f**, Median DMR and MSE for TranScouter and delta-average in low-sign-match target-pair subsets. **g**, Importance of condition-aware pairing and additional architectural components. Normalized MSE is defined as model MSE divided by the MSE of a control-expression predictor, so values near 1 indicate control-level performance.

#### Dependence on similarity between test and training conditions

We next quantified how predictive difficulty scales with the distance between a held-out condition and those observed during training. For a held-out condition *c^∗^*, we defined this distance as the minimum Euclidean distance between the encoded control representation of *c^∗^* and any training condition:

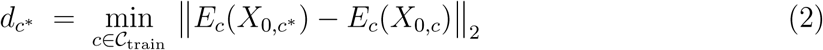

where *C*_train_ denotes the training conditions and *E_c_* is the control–cell encoder of TranScouter. As shown in Figure 4b, larger *d_c_∗* was associated with higher error, with a statistically significant yet modest positive correlation (Pearson *r* = 0.22, *p <* 0.001). This relationship suggests that the condition embedding space captures biologically meaningful structure: conditions that are closer in this learned representation tend to transfer better. At the same time, the modest correlation indicates that useful robustness can be retained even when the target condition is not especially close to training conditions. We also observed that variance in MSE increased with distance, consistent with the possibility that more distinct biological conditions introduce interaction patterns that are less well represented during training. Together with the training-diversity analysis, these results support a representation-based view of cross-condition prediction: performance is associated with both the breadth of condition-space coverage in training and the proximity of the held-out condition to that covered space.

### Perturbation-effect transferability

The preceding analyses focus on the condition axis. We next asked a complementary question about the perturbation axis: to what extent can the response to a perturbation in a held-out condition be approximated by transferring the perturbation-induced change observed in other conditions? This question is motivated in part by recent cross-condition models such as TxPert [24], whose cross-cell-line formulation can make effective use of a perturbation-specific expression shift added to a target-condition control state. Our goal here is not to approximate TxPert, whose perturbation effect is learned from graph-based perturbation representations, but to isolate the simpler empirical structure underlying this class of transfer: whether the average observed effect of the same perturbation across other conditions is already informative for the target condition.

To test this, we defined a delta-average diagnostic baseline. Let *µ*(*c, p*) denote the mean expression profile for perturbation *p* in condition *c*, and let *µ*(*c,* ctrl) denote the corresponding control profile. For a target held-out condition *c^∗^*, we define the perturbation-induced change observed in a training condition *c* as

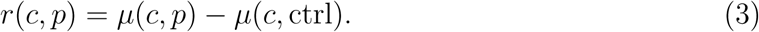

If perturbation *p* is observed in a set of non-held-out training conditions 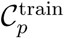, the deltaaverage prediction is

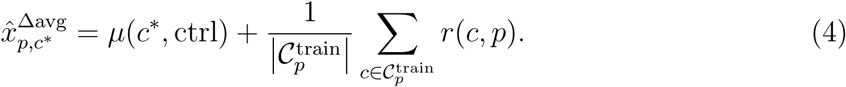

Thus, delta-average separates the prediction into a target-condition baseline component and an empirically transferred perturbation-effect component. This differs from the averaging baseline defined in Equation 1, which directly averages perturbed expression profiles and therefore does not explicitly preserve the baseline state of the target condition. Delta-average is defined only when the target perturbation has been observed in at least one non-held-out training condition; when applied to the strict unseen-perturbation scenario, it should therefore be interpreted as an optimistic diagnostic rather than as a deployable prediction method.

On Jiang24, delta-average performed strongly by directional accuracy (Figure 4c). Its DMR was lower than TranScouter in both the seen- and unseen-perturbation scenarios, with the unseen comparison benefiting from the same optimistic access to target-perturbation responses described above. The MSE comparison was less uniformly favorable to delta-average: TranScouter retained lower MSE in the seen-perturbation scenario, while the two methods were closer in the optimistic unseen comparison. Thus, the aggregate Jiang24 results raised a more specific question: whether the DMR advantage of delta-average reflected a uniform weakness of TranScouter across target pairs, or whether the two approaches differed across subsets with different levels of perturbation-effect transferability.

To examine this heterogeneity, we stratified target perturbation-condition pairs by the directional transferability of the average perturbation effect. For each target pair (*c^∗^, p*), we computed a sign-match score on the same DEG set used for DMR. This score is equivalent to one minus the DMR of the delta-average prediction for that target pair: a high sign-match means that the average response change transferred from other conditions agrees in sign with the true target response for most DEGs, whereas a low sign-match means that this transfer shortcut is directionally unreliable. Across Jiang24 target pairs, the median sign-match was 0.92 (Figure 4d), corresponding to a median delta-average DMR of 0.08. Thus, many Jiang24 target pairs contain a highly transferable directional component, making them favorable cases for an empirical delta-transfer diagnostic.

We next asked whether the DMR advantage of delta-average was distributed uniformly across target pairs or concentrated in cases where this directional transfer shortcut was strong. Sign-match was strongly associated with the DMR advantage of delta-average relative to TranScouter, defined as DMR_TranScouter_ − DMR_Δavg_ (Spearman *ρ* = 0.58; Figure 4e). This association indicates that the relative advantage of delta-average was largest when the average perturbation-induced change from other conditions already matched the target response direction. Conversely, when sign-match was low, the relationship reversed. Among target pairs with sign-match ≤ 0.55 (112/1236 target pairs; 9.06%), TranScouter achieved lower median DMR (0.25 versus 0.56) and lower median MSE (0.10 versus 0.21) than delta-average (Figure 4f). The same pattern persisted under a broader threshold of sign-match ≤ 0.75 (282/1236 target pairs; 22.82%), where TranScouter again showed lower median DMR (0.20 versus 0.38) and lower median MSE (0.10 versus 0.20). These results suggest that TranScouter is not uniformly weaker than delta-average on Jiang24; rather, delta-average is strongest in the many target pairs where directional perturbation effects are readily transferable, whereas TranScouter performs better in the subset where this transfer shortcut is unreliable.

High sign-match does not imply that the full perturbation-response profile is condition-invariant. Sign-match only evaluates whether the averaged response change has the correct direction for each DEG; it does not evaluate relative magnitudes, gene-wise ranking, or the continuous structure of the response vector. To examine this distinction, we computed a train-target response-correlation score for each target pair. For each training condition *c* and perturbation *p*, we correlated the response-change vector in condition *c* with the response-change vector in the held-out condition *c* on the target DEG set *D_p,c_∗* :

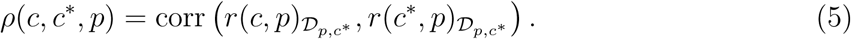

We then averaged this quantity across training conditions:

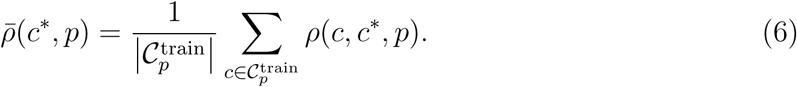

Across Jiang24 target pairs, the median *ρ̄*(*c^∗^, p*) was only 0.08, despite the median sign-match of 0.92. This indicates that Jiang24 contains a conserved directional component that delta-average can exploit, but that individual response profiles across conditions remain weakly correlated. In this sense, the strong directional performance of delta-average should not be interpreted as evidence that perturbation responses are fully condition-independent; rather, it shows that directional conservation and continuous response-profile similarity can be sub-stantially different aspects of cross-condition transfer.

Taken together, these analyses indicate that the Jiang24 benchmark contains many target pairs for which perturbation-induced directions are highly transferable across biological conditions, explaining why delta-average performs strongly on DMR. This is biologically plausible: perturbations affecting core signaling components, broad stress responses, or shared cellular programs can induce partially conserved transcriptional directions across contexts. However, the value of multi-condition perturbation experiments lies not only in recovering these shared components, but also in identifying where perturbation responses deviate from them. Such deviations can reflect condition-dependent biology, including differences in pathway activity, stimulation state, or cell-line background.

Consistent with this view, the transfer shortcut was not universal. Jiang24 contained a nonnegligible low-sign-match subset in which TranScouter performed better, indicating cases where the average perturbation effect from other conditions was directionally unreliable. Moreover, in McFaline [43], a secondary multi-condition perturbation dataset, the same diagnostic showed much weaker directional transferability, with median sign-match close to random directional agreement and correspondingly weaker delta-average performance (Supplementary Note 4). Thus, delta-average is best interpreted as a diagnostic for transferable perturbation-effect structure rather than as a general replacement for cross-condition modeling. More broadly, these results support a mixed view of cross-condition perturbation prediction: some target pairs are dominated by conserved perturbation-effect directions that can be captured by simple transfer, whereas others require modeling condition-dependent deviations in direction, magnitude, or gene-wise response structure.

### Extension from single-condition to cross-condition

Finally, we examined modeling choices required to use condition information correctly in cross-condition prediction, using the relationship between Scouter and TranScouter as an illustrative example. In Scouter, all cells are assumed to be drawn from the same biological condition, so matching control and perturbed cells does not require an explicit condition constraint. In TranScouter, by contrast, each perturbed cell is paired only with control cells from the same biological condition, so that the model learns perturbation response relative to the appropriate condition-specific baseline. Once multiple biological conditions are present, this alignment becomes essential: otherwise the model no longer receives a coherent relationship between baseline state and perturbation response. To test this directly, we evaluated a TranScouter variant without condition-aware pairing, such that a control cell from one biological condition could be matched with a perturbed cell from another condition. For this ablation, we report normalized MSE, defined as model MSE divided by the MSE from a naive predictor using average control-cell expression, as a reference point for when a method ceases to extract meaningful perturbation-specific signal. Under this setting, as shown in Figure 4g, normalized MSE approached 1, indicating performance close to the control predictor. This result suggests that preserving the alignment between perturbation response and the biological condition in which it is measured is a basic requirement for cross-condition prediction rather than merely an implementation detail.

We next asked whether the additional architectural components introduced in TranScouter beyond this condition-aware pairing provide further benefit. To test this, we adapted Scouter to the cross-condition setting by introducing condition-aware pairing. This variant is otherwise similar to TranScouter, except that it lacks the gene-embedding encoder and the reintroduction of the condition embedding at the bottleneck. Relative to the full TranScouter model, this simplified variant showed a modest increase in MSE of approximately 15%, as demonstrated in Figure 4g. Taken together, these analyses suggest that the essential modeling requirement is to preserve the correct alignment between perturbation response and biological condition, while additional architectural refinements can provide further, but comparatively smaller, gains. This supports the view that effective cross-condition prediction depends less on elaborate architecture alone than on representing and aligning the perturbation and condition axes appropriately.

## 3 Discussion

In this work, we studied cross-condition prediction of transcriptional responses to gene perturbations as a structured problem with distinct information scenarios. Specifically, we considered both a seen-perturbation scenario, where the target perturbation has been observed in other biological conditions, and an unseen-perturbation scenario, where the target perturbation is absent from all training perturbation-condition pairs. Evaluating both scenarios within the same benchmark framework shows that whether the target perturbation has been observed under other conditions strongly shapes what kind of generalization is being tested. TranScouter provides a lightweight representation-based instantiation for studying this problem, and the accompanying analyses characterize condition-space coverage, perturbation-effect transferability, and condition-aware modeling choices as factors associated with cross-condition performance.

A central implication of this study is that cross-condition prediction can be approached through paired representations of perturbation identity and target biological condition. Prior work such as Scouter and GenePert showed that perturbation extrapolation can be supported by informative continuous representations of perturbed genes, including text-derived embeddings. Here, we apply the same representational logic to the condition axis by using control-cell transcriptomic profiles from the target condition as a representation of biological state. In this formulation, the control transcriptome serves both as the baseline on which perturbation effects are expressed and as a coordinate for relating held-out conditions to conditions observed during training. The competitive performance of a relatively simple encoder-decoder model across both scenarios suggests that effective cross-condition prediction does not necessarily require elaborate architecture as a starting point; it depends on whether perturbation and condition are represented in forms that allow meaningful transfer and whether those representations are correctly aligned during training.

The delta-average and sign-match analyses reveal a structural feature of cross-condition perturbation prediction: benchmark performance reflects a mixture of conserved perturbation effects and condition-dependent deviations. On Jiang24, many perturbation-condition pairs contain a conserved directional response component, allowing a simple empirical transfer of average perturbation-induced changes to perform strongly by DMR. This should not be viewed as a failure of the benchmark or as evidence that cross-condition modeling is unnecessary; rather, it reflects a biologically plausible structure in which perturbations affecting shared pathways can induce partially conserved transcriptional directions across conditions. At the same time, the non-negligible low-sign-match subset shows that this transfer shortcut is not universal, especially in the McFaline analysis (Supplementary Note 4). The cases in which perturbation responses deviate from the conserved average effect are precisely where multi-condition perturbation experiments become especially informative, because such deviations can reveal condition-dependent pathway activity, stimulation effects, or cell-line-specific response programs. This suggests that future cross-condition benchmarks may benefit from reporting not only aggregate prediction accuracy, but also the transferability structure of the target pairs being evaluated.

At the same time, our analyses also reveal limitations of the current model and evaluation setting, pointing to directions for future work. Predictive accuracy declined when the training set provided poor coverage of the biological condition space: performance dropped sharply when training diversity was reduced to only a few conditions, and error increased when the test condition was distant from those represented in training. These observations underscore the inherent difficulty of extrapolating across poorly represented regions of biological space and point to the need for broader and more diverse training collections to ensure reliable cross-condition generalization.

The main evaluation in this study is centered on Jiang24, which spans thirty biological conditions and provides a relatively broad condition space for the present study of genetic perturbation responses. Nevertheless, broader coverage across more cell types, treatments, and cellular states would further strengthen evaluation of cross-condition generalization. A straightforward way to expand condition coverage would be to combine perturb-seq datasets from multiple studies. However, in this task, the relevant unit of evaluation is not simply the number of dataset files, but the coverage and coherence of the perturbation-by-condition space. Multi-study integration introduces practical challenges: study-specific batch effects can be large, the set of overlapping genes can shrink substantially, and dataset identity may become confounded with the biological conditions being evaluated. These issues are especially problematic in the cross-condition setting, where the signal of interest is precisely variation across biological conditions, with a more detailed rationale provided in Supplementary Note 5. We therefore prioritize Jiang24 for the main analysis because it provides a relatively large set of internally consistent biological conditions, and include McFaline [43], a smaller multi-condition dataset spanning fifteen biological conditions, as additional analysis in Supplementary Note 6. As larger harmonized perturbation atlases spanning more biological conditions become available, they will enable broader and more definitive evaluation of this task.

Taken together, this study argues for treating cross-condition perturbation prediction as a structured modeling and evaluation problem rather than as a single aggregate benchmark. Separating seen- and unseen-perturbation scenarios clarifies what information is available for transfer, while the delta-average and sign-match diagnostics help distinguish target pairs dominated by conserved perturbation effects from those shaped by condition-dependent deviations. TranScouter provides one lightweight representation-based approach for studying this setting, showing that perturbation and condition representations can be combined effectively without relying on highly elaborate architecture.

Looking ahead, the limitations of the present evaluation are likely to be alleviated as increasingly large and diverse perturbation compendia become available. For example, although Tahoe100M [44] is a drug-perturbation resource and therefore not suitable for the genetic-perturbation task studied here, it nevertheless exemplifies the scale and diversity of perturbation atlases that are already becoming achievable. Similar resources for genetic perturbations would enable more systematic evaluation of condition representations, more reliable characterization of perturbation-effect transferability, and stronger pretrained models for cross-condition generalization. In this sense, TranScouter should be viewed less as an endpoint than as a practical starting point for studying how perturbation and condition information can be combined in increasingly diverse perturbation-response datasets.

## 4 Methods

### 4.1 Problem formulation

We formulate cross-condition perturbation prediction as the task of predicting the transcriptional response to a gene perturbation in a biological condition that is entirely absent during training. Let *P* denote the set of perturbation targets and *C* the set of biological conditions. In this study, a biological condition refers to the baseline cellular context in which a perturbation is applied, such as a cell line, stimulation state, treatment condition, or their combination. For a perturbation target *p* ∈ *P* and condition *c* ∈ *C*, let *x_p,c_* ∈ R*^G^* denote the post-perturbation expression profile over *G* genes, and let *x*_0_*_,c_* ∈ R*^G^* denote the corresponding control expression profile for unperturbed cells in condition *c*.

The goal is to learn a function

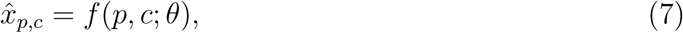

where *x*^*_p,c_* is the predicted post-perturbation expression profile and *θ* denotes learnable model parameters. At this level of formulation, *p* and *c* denote the perturbation target and biological condition, respectively; their numerical representations are specified in later sections. Thus, the model is asked to predict the expression profile that would be observed if perturbation *p* were applied under condition *c*.

We evaluate this task under condition-level generalization. During training, both control and perturbed expression profiles are observed only for a set of training conditions *C*_train_, while evaluation is performed on held-out conditions *C*_test_ with *C*_train_ ∩ *C*_test_ = ∅□. We distinguish two scenarios according to whether the perturbation target has been observed during training. In the seen-perturbation scenario, the test perturbation *p* has been observed in at least one training condition, but the pair (*p, c*) is unobserved because *c* ∈ *C*_test_. In the unseen-perturbation scenario, *p* is absent from all training perturbation-condition pairs, so prediction requires extrapolation along both the perturbation and condition axes. In both scenarios, control cells from the held-out condition are available at evaluation time as the representation of the target biological condition.

### 4.2 Datasets

We used Jiang24 [33] as the primary benchmark for cross-condition perturbation prediction. Jiang24 is a large-scale CRISPRi Perturb-seq dataset designed to profile signalling-regulator perturbations across multiple cellular and environmental contexts. The dataset contains six cancer cell lines (A549, BxPC-3, HAP1, HT-29, K562, and MCF-7), each exposed to one of five cytokine or stimulation conditions (IFNB, IFNG, INS, TGFB, and TNFA), yielding 30 biological conditions in total. Control cells are available for all 30 conditions. After preprocessing and filtering as described below, the dataset used in this study contains 218 non-control perturbation targets and 1,656 observed perturbation-condition pairs.

Jiang24 is particularly suitable for this study because it provides a relatively broad set of biological conditions measured within a coherent experimental framework. The cell line and stimulation axes introduce substantial variation in baseline transcriptional state, making the dataset appropriate for evaluating whether a model can transfer perturbation-response information to held-out conditions. We visualize the control-cell transcriptomes across the 30 conditions in Supplementary Note 1, which illustrates that these conditions occupy distinct regions of transcriptional space. Because perturbation coverage is not complete across all conditions, the dataset also naturally reflects the sparse perturbation-by-condition structure that motivates cross-condition prediction. We summarize this structure using a perturbation-by-condition availability heatmap and perturbation-coverage statistics in Supplementary Note 1 as well.

We additionally evaluated TranScouter on McFaline [43] as a supplementary dataset. McFaline is a sci-Plex-GxE chemical-genetic CRISPRi screen of kinase perturbations across three glioblastoma cell lines and five treatment settings, yielding 15 biological conditions. In our processed version, the dataset contains 519 non-control perturbation targets. McFaline is smaller, substantially sparser, and required dataset-specific preprocessing, and we use it as an additional analysis; details of its preprocessing and evaluation are provided in Supplementary Note 6.

### 4.3 Data preprocessing

For Jiang24, we started from the five treatment-specific Seurat [45] objects released by the original study and converted them to AnnData format [46]. We then followed the PerturBench quality-control workflow [47] to merge the treatment-specific objects, standardize metadata, and perform cell-level and gene-level filtering. Starting from this curated cell-level dataset, we performed preprocessing to construct the benchmark analyzed here.

A key preprocessing choice was the level at which single-cell profiles were aggregated. Existing perturbation-response studies have used both single-cell-level training [19, 48, 49] and fully collapsed pseudobulk profiles [17, 18, 20]. In our setting, both extremes raise concerns. Retaining individual cells preserves cell-level heterogeneity, but introduces substantial computational overhead and requires pairing perturbed cells with control cells despite the absence of a natural one-to-one correspondence. Conversely, collapsing all cells from the same perturbation-condition pair into a single profile discards sample and batch structure, and leaves too few replicates for robust differential expression analysis.

We therefore used an intermediate replicate-level aggregation strategy. Single-cell counts were summed within each unique combination of cell line, treatment, perturbation target, sample, and batch. Each aggregate profile therefore represents one perturbation or control group measured within a specific biological condition and experimental stratum, rather than a single cell or a fully collapsed perturbation-condition average. This strategy reduces singlecell noise while preserving replicate structure for model training and DEG identification. In Jiang24, each observed perturbation-condition pair was represented by a median of 32.0 aggregate profiles and a mean of 31.3 aggregate profiles.

After aggregation, we normalized the aggregate count profiles using `scanpy.pp.normalize_total` and applied log transformation with `scanpy.pp.log1p`. We then selected 5,000 highly variable genes using `scanpy.pp.highly variable genes` with the biological-condition label as the batch key, `flavor=“seurat_v3“`, and `n_top_genes=5000`. The resulting processed Jiang24 dataset was used as the common benchmark for training and evaluating all methods unless otherwise specified.

Evaluation metrics were computed on differentially expressed genes (DEGs). For each biological condition, each perturbation group was compared against the corresponding control group using `scanpy.tl.rank_genes_groups` with `method=“wilcoxon“` and `rankby_abs=True`. Because large single-cell datasets can yield statistically significant but very small expression differences, we required both a p-value cutoff of 0.1 and an absolute log fold-change threshold of 1.5. This effect-size filter was used to focus evaluation on genes with appreciable perturbation-associated changes. For each perturbation-condition pair (*p, c*), we retained up to 25 DEGs, denoted *D_p,c_*, for evaluation. These DEG sets were used only for computing evaluation metrics.

### 4.4 Train-validation-test splitting

We evaluated cross-condition generalization using 10 condition-level splits of Jiang24. In each split, 3 biological conditions were held out for testing, 2 additional biological conditions were held out for validation, and the remaining 25 biological conditions were used for training. The split was performed at the level of biological conditions, so for a held-out test condition, neither its control profiles nor its perturbed profiles were used during model training. At evaluation time, however, control profiles from the held-out condition were provided as the condition representation. This mirrors the intended use case in which a new biological condition can be profiled in an unperturbed control state, while perturbation experiments in that condition are unavailable. Thus, access to target-condition controls is part of the task definition rather than information leakage from the held-out perturbation responses.

Within each split, we further masked a subset of perturbation targets to evaluate extrapolation across perturbations. Specifically, approximately 10% of perturbation targets appearing in the validation or test conditions were removed from the training set across all training conditions. Test cases in held-out biological conditions were then separated into two scenarios. In the seen-perturbation scenario, the perturbation target was observed in at least one training condition but not in the held-out target condition. In the unseen-perturbation scenario, the perturbation target was absent from all training perturbation-condition pairs. Main performance summaries were computed on held-out biological conditions and aggregated across the 10 splits.

### 4.5 Perturbation and condition representations

TranScouter represents each prediction input using two complementary components: a representation of the perturbation target and a representation of the target biological condition. For perturbation targets, we used pretrained text-derived gene embeddings. Rather than introducing a new gene-embedding method, we treated the embedding source as a model choice and considered three existing embedding sets: GenePT [27] embeddings generated from NCBI gene summaries, an expanded GenePT variant that additionally incorporates UniProt descriptions when available, and scELMo [50] gene embeddings, which use LLM-generated descriptions of gene function to represent perturbation targets in single-cell modeling tasks. Gene symbols in the perturbation dataset were matched to the corresponding embedding entries before model training.

Biological conditions were represented using control-state transcriptomic profiles. For a target condition *c*, the condition input is the expression profile *x*_0_*_,c_* of unperturbed cells measured under the same cell line and treatment context. This representation is intended to capture the baseline transcriptional state on which the perturbation acts, including cellline identity, stimulation state, and other condition-specific expression programs. In the processed Jiang24 benchmark, both the condition representation and the predicted postperturbation profile are defined over the same 5,000 highly variable genes.

### 4.6 Model architecture

TranScouter is implemented as a lightweight encoder-decoder model with separate branches for perturbation and condition inputs. Given a perturbation target *p*, represented by a pretrained gene embedding *e_p_*, and a matched control expression profile *x*_0_*_,c_* from the target biological condition, the model predicts the corresponding post-perturbation expression profile *x*^*_p,c_*. The pretrained gene embedding matrix is kept fixed during model training; only the downstream neural network parameters are learned.

The perturbation embedding and control expression profile are first encoded by separate multilayer perceptrons (MLP). Let *E_p_* denote the perturbation encoder and *E_c_* denote the condition encoder:

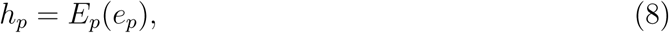

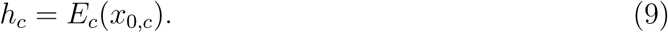

The two encoded representations are concatenated and projected into a bottleneck latent representation,

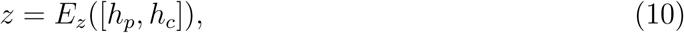

where [·, ·] denotes concatenation. Although the implementation supports variational sampling from this bottleneck, all main experiments used the deterministic latent representation.

To preserve condition-specific information during reconstruction, TranScouter reintroduces the encoded condition representation before decoding. Specifically, the decoder receives the concatenation [*z, h_c_*] and outputs the predicted post-perturbation expression profile:

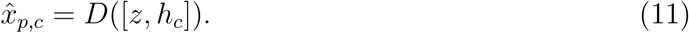

The output is a 5,000-dimensional expression vector over the highly variable genes used in the processed benchmark dataset. This design allows the model to combine perturbation-specific information from the gene embedding with condition-specific information from the matched control state while making no assumption that perturbation effects are identical across biological conditions.

### 4.7 Training objective and optimization

TranScouter was trained to reconstruct the observed post-perturbation expression profile from the perturbation embedding and matched control-state expression profile. We adopted the autofocus direction-aware loss from GEARS [17]. This loss contains an autofocus reconstruction term,

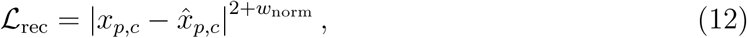

where *w*_norm_ controls the degree to which larger expression errors are upweighted. The loss function also includes a direction-aware term comparing the predicted and observed signs of expression change relative to the matched control profile:

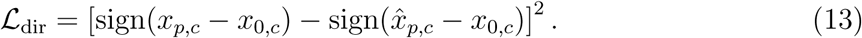

The total training loss was

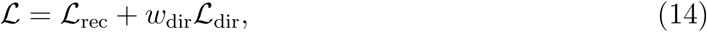

averaged over genes and perturbation-condition groups within each mini-batch. Although the implementation supports an optional variational bottleneck with a KL-divergence term, all main experiments used deterministic encoding and therefore did not include this term.

Models were optimized with Adam using mini-batch training and exponential learning-rate decay. We used gradient clipping with maximum norm 1.0 and early stopping based on validation loss with patience of 5 epochs. The perturbation embedding source, layer dimensions of *E_p_* and *E_c_*, dropout rate, batch size, learning rate, learning-rate decay factor, reconstruction-loss exponent *w*_norm_, and direction-loss weight *w*_dir_ were treated as tunable hyperparameters and selected based on validation loss.

### 4.8 Comparator methods

We compared TranScouter with three existing perturbation-response methods that could be adapted to parts of the cross-condition prediction task: scGen [22], trVAE [23], and CellOracle [37]. Because these methods were originally developed for related but not identical endpoints, we adapted each method according to its official usage pattern and evaluated it only in the scenario where its required inputs and assumptions were applicable.

For scGen, we followed the official perturbation-prediction tutorial (https://scgen.readthedocs.io/en/stable/tutorials/scgen_perturbation_prediction.html). scGen was originally designed to transfer a learned perturbation effect to a held-out cell type or context by estimating a latent-space shift between control and perturbed states. To adapt this framework to our setting, we used perturbation identity as the transferable state axis and biological condition as the context label. In the scGen implementation, this corresponds to setting batch key to the field encoding perturbation identity and labels key to the field encoding biological condition. For each held-out biological condition and seen perturbation, ctrl key was set to the control state, stim key to the target perturbation, and celltype to predict to the held-out biological condition. Thus, scGen estimated the perturbation-associated latent shift from training conditions and applied that shift to control profiles from the held-out condition. Since scGen requires the target perturbation state to be observed during training in order to estimate this shift, we evaluated scGen only in the seen-perturbation setting.

For trVAE, we followed the perturbation-transfer examples provided with the official implementation (https://github.com/theislab/trvaep/tree/master/example). trVAE is a conditional variational autoencoder that predicts a target state by encoding source profiles and decoding them under a specified target condition label. To adapt trVAE to cross-condition perturbation prediction, we used perturbation identity as the conditional label. In the trVAE implementation, condition key was set to the field encoding perturbation identity. For each held-out biological condition and seen perturbation, the source matrix x consisted of control profiles from the held-out condition, the source labels y indicated the control state, and target indicated the perturbation to be predicted. The model therefore encoded target-condition control profiles and decoded them under the target perturbation label. As with scGen, trVAE was evaluated only for seen perturbations, because the target perturbation label must be present during training.

For CellOracle, we followed the official tutorials for base-GRN preparation, GRN construction, and in silico TF perturbation simulation (https://morris-lab.github.io/CellOracle.documentation/tutorials/). CellOracle is a GRN-based framework that infers condition-specific regulatory networks and simulates how transcription-factor perturbations propagate through those networks. For each held-out biological condition, we constructed a CellOracle model using control profiles from that condition and the built-in human promoter base GRN. Following the tutorial workflow, control profiles were imported into an Oracle object, regulatory links were inferred and filtered, cluster-specific TF dictionaries were extracted, and the GRN was fitted for simulation. We then simulated knockout-like perturbation of each eligible target gene by setting its expression to zero and propagating the perturbation through the fitted GRN. Because our benchmark evaluates post-perturbation expression profiles rather than CellOracle’s usual vector-field or transition outputs, we extracted the simulated expression layer before downstream transition or embedding-shift analysis and used it as the predicted expression profile. CellOracle was evaluated only for perturbations whose target genes were present among the active regulatory genes in the CellOracle model, and therefore its metrics were computed on a TF/regulatory-gene subset of the unseen-perturbation test cases rather than on the full unseen-perturbation benchmark.

### 4.9 Baseline methods

We also included simple reference predictors to contextualize model performance. The perturbation-averaging baseline uses the same technical procedure in both seen- and unseen-perturbation analyses: for a target perturbation *p* in a held-out condition *c*, it predicts the response by averaging observed expression profiles for the same perturbation *p* across non-test biological conditions. This baseline tests whether a condition-independent average perturbed profile is sufficient, and therefore provides a direct reference for the value of condition-specific modeling. In the seen-perturbation setting, this is a natural baseline because those same-perturbation profiles are part of the training data. In the unseen-perturbation setting, however, the same calculation should be interpreted differently: by construction, perturbation *p* is masked from all training conditions for TranScouter and CellOracle, whereas the averaging baseline is computed from the masked same-perturbation profiles in other biological conditions. It therefore has access to perturbation-specific information deliberately withheld from TranScouter and should be viewed as an optimistic reference rather than as a deployable predictor under the strict unseen-perturbation task definition.

For the unseen-perturbation setting, we additionally implemented a correlation-based heuristic inspired by CellPB [38]. For each split, we computed a gene-gene Pearson correlation matrix *R* from training perturbed profiles. To predict knockdown of target gene *p* in condition *c*, the predicted expression of gene *g* was computed as

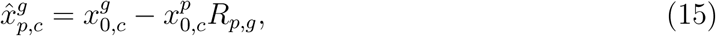

where 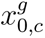 is the control expression of gene *g* in condition *c*, 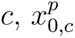 is the control expression of the target gene, and *R_p,g_* is the Pearson correlation between genes *p* and *g* estimated from the training data. Genes positively correlated with the target gene are therefore shifted downward after simulated knockdown, whereas negatively correlated genes are shifted upward.

### 4.10 Evaluation metrics

Evaluation was performed for each perturbation-condition pair using the corresponding DEG set *D_p,c_* defined above. Let 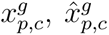, and 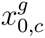 denote the observed, predicted, and control expressions of gene *g* under perturbation *p* and condition *c*, respectively. Prediction error was measured by mean squared error (MSE) over DEGs:

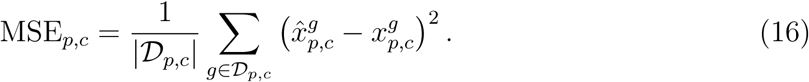

Lower MSE indicates closer agreement between predicted and observed post-perturbation expression.

To evaluate whether the predicted response recovered the direction of perturbation-induced changes, we computed directional mismatch rate (DMR):

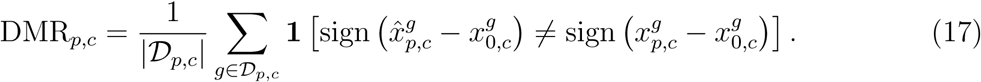

DMR ranges from 0 to 1, with lower values indicating better agreement between predicted and observed directions of change. Pearson correlation coefficient (PCC) was also computed on DEG-level predictions as a supplementary diagnostic when defined.

Metrics were first computed for individual perturbation-condition pairs. Unless otherwise specified, condition-level summaries were obtained by taking the median across perturbations within each biological condition to reduce sensitivity to outlier perturbations.

### 4.11 Cross-condition generalization analyses

We performed additional analyses on Jiang24 to characterize factors associated with cross-condition generalization. To assess the effect of training-condition diversity, separate condition-level splits were generated where we varied the number of biological conditions available for training. TranScouter was trained with 5, 10, 15, 20, or 25 training conditions and evaluated on held-out biological conditions. This analysis was used to test how much diversity across biological contexts is needed before cross-condition prediction becomes stable.

To examine whether performance depends on similarity between training and test conditions, we used the learned condition encoder *E_c_* to embed control profiles from both training and held-out biological conditions. For each held-out condition *c^∗^*, we computed its distance to the training set as defined in Eq.2. We then compared this distance with prediction error on the corresponding held-out condition and quantified the association using Pearson correlation.

The delta-average diagnostic and sign-match score were defined as described in the Results. For each target pair, all quantities were computed using only non-held-out biological conditions in which the same perturbation was observed, and evaluation used the same DEG sets and MSE/DMR definitions as the main benchmark. When reported for the unseen-perturbation scenario, delta-average was treated as an optimistic diagnostic because it uses perturbation-specific response information withheld from TranScouter under the task definition.

Low-sign-match subsets were defined by thresholding the target-pair-level sign-match score. For the train-target response-correlation analysis, response-change correlations were computed on the target DEG set by correlating each training-condition response-change vector with the target-condition response-change vector and averaging across available training conditions. The same diagnostic procedure was applied to McFaline for the supplementary transferability analysis (Supplementary Note 4).

To test the importance of condition-aware pairing, we constructed an ablation in which control profiles were randomly permuted before being paired with perturbed profiles during dataset construction. This preserves the same expression profiles and split structure, but breaks the alignment between each perturbation response and the biological condition in which that response was measured. The resulting model was trained and evaluated using the same protocol as the full model.

Finally, we evaluated an architecture ablation where condition-aware pairing was retained but the perturbation-embedding encoder and the reintroduction of the condition embedding at the decoder bottleneck were removed. This comparison was used to assess the contribution of the additional TranScouter architecture components after the basic requirement of correct condition-response pairing had been satisfied.

### 4.12 Computing resources

Runtime was measured under the same hardware setting for each compared method. CPU runtime was measured on a MacBook Pro (2022) with an Apple M2 eight-core CPU and 16 GB of RAM. GPU runtime was measured on a server with a 32-core Intel Xeon Gold 6326 CPU, 256 GB of RAM, and an NVIDIA A40 GPU with 48 GB of GPU memory. Each reported runtime reflects model training across all 10 data splits.

### 4.13 Data availability

The Jiang24 dataset used for the main analyses was obtained from the processed data released by the original study at Zenodo (https://zenodo.org/records/14518762); raw sequencing data for that study are available from GEO under accession code GSE281048. The McFaline dataset used for supplementary analyses was obtained from GEO under accession code GSE225775, where the original study made processed and raw data available. Both GenePT embedding versions were downloaded from Zenodo (https://zenodo.org/records/10833191), and scELMo embeddings were downloaded from the scELMo resource site (https://sites.google.com/yale.edu/scelmolib).

### 4.14 Code availability

TranScouter is available as a Python package at https://github.com/PancakeZoy/TranScouter. Code for reproducing the analyses in this study will be made available at https://github.com/PancakeZoy/TranScouter_misc. The reproducibility repository will include documented scripts for obtaining data, preprocessing, training models, generating predictions, evaluating performance, and conducting method comparisons.

