## Supplementary Material for "A structured study of cross-condition prediction of transcriptional responses to gene perturbations"

### Supplementary Materials

#### Supplementary Note 1: Jiang24 condition and perturbation coverage

To further characterize the structure of the Jiang24 benchmark, we examined both the transcriptional separation among biological conditions and the availability of perturbations across those conditions, as shown in Supplementary Figure 1. This analysis serves two purposes. First, it verifies that the cell line-treatment combinations used as biological conditions correspond to distinct baseline transcriptional states. Second, it shows that perturbation coverage is sparse across conditions, which is precisely the setting that motivates cross-condition prediction.

For the condition-level visualization, we embedded 84,269 unperturbed control cells before aggregation and colored cells by their cell line-treatment condition. The resulting UMAP shows that many biological conditions occupy distinct regions of transcriptional space, with separation driven by both cell-line identity and stimulation state. This supports the view that the held-out conditions in our benchmark represent meaningful changes in baseline biological context rather than near-duplicate samples. Accordingly, predicting perturbation responses across these conditions requires a model to account for variation in the underlying cellular state.

We also summarized perturbation availability using a perturbation-by-condition heatmap. The Jiang24 benchmark contains 219 perturbation labels including the control label, corresponding to 218 non-control perturbation targets. These perturbations are not uniformly observed across all 30 biological conditions. After preprocessing and filtering, the benchmark used in this study contains 1,656 observed non-control perturbation-condition pairs out of 6,540 possible pairs. Thus, the dataset contains a broad union of perturbations across conditions, but each condition contains only a subset of the full perturbation set.

Together, these two views illustrate why Jiang24 is well suited for evaluating cross-condition perturbation prediction. The control-cell UMAP shows that the biological conditions differ appreciably in baseline transcriptional state, while the coverage heatmap shows that the perturbation-by-condition matrix is incomplete. This combination creates a realistic benchmark in which models must transfer information across biological contexts rather than simply interpolate among fully observed perturbation-condition pairs.

#### Supplementary Note 2: PCC analysis

We reported Pearson correlation coefficient (PCC) as a supplementary diagnostic metric to assess whether predicted and observed DEG-level response profiles had similar overall patterns. PCC is complementary to the primary metrics used in the main text, but it should be interpreted differently from MSE and DMR. Because PCC is invariant to shifts and rescaling of the predicted vector, it can remain high even when absolute expression values are poorly estimated or when individual DEG directions are not consistently recovered. We therefore use PCC to summarize response-pattern similarity, while retaining MSE and DMR as the primary measures of prediction accuracy and directional correctness.

PCC was computed over the same DEG sets used for MSE and DMR, with one additional stability criterion: PCC was evaluated only for perturbation-condition pairs with more than ten DEGs. This avoids over-interpreting correlations computed from very short DEG vectors. Condition-level summaries were then computed as in the main evaluation.

In the seen-perturbation setting, PCC was broadly consistent with the main performance results, as illustrated in Supplementary Figure 2. Aggregated across biological conditions, the median PCC was 0.72 for TranScouter, compared with 0.63 for scGen, 0.65 for trVAE, and 0.22 for the averaging baseline. Thus, TranScouter showed the strongest overall response-pattern similarity among the evaluated methods in the seen-perturbation regime.

In the unseen-perturbation setting, TranScouter also achieved high PCC across the full unseen-perturbation benchmark, with an aggregated median PCC of 0.70, compared with 0.54 for the correlation-based reference and 0.19 for the averaging baseline, as illustrated in Supplementary Figure 3. CellOracle achieved a higher aggregated median PCC of 0.85 where it was applicable. However, this value was computed from only 20 perturbation-condition pairs involving TF perturbations compatible with CellOracle's GRN-based simulation framework, whereas TranScouter, the correlation-based reference, and the averaging baseline were evaluated on the full unseen-perturbation test set of 72 perturbation-condition pairs. The missing CellOracle bars in Supplementary Figure 3 indicate held-out conditions for which CellOracle could not be evaluated because the relevant perturbations were outside this TF-based scope.

Together, these results show that PCC provides a useful complementary view of response-pattern similarity, while reinforcing the need to interpret correlation-based summaries alongside metrics that directly quantify absolute error and directional correctness.

#### Supplementary Note 3: Additional qualitative biological examples

We examined two additional Jiang24 perturbation-condition pairs to illustrate how DEG-level directional errors affect biological interpretation. These examples complement the TYK2 and TNFR1 cases discussed in the main text and were selected because the perturbed genes encode receptor components directly relevant to the cytokine signaling contexts in which they were evaluated.

In the seen-perturbation setting, we examined IFNAR2 perturbation in A549 cells under IFN-$\beta$ stimulation (Supplementary Figure 4). IFNAR2 is a receptor subunit required for type I interferon signaling, and IFN-$\beta$ stimulation induces a broad interferon-stimulated gene program through this pathway. Therefore, IFNAR2 knockdown is expected to attenuate induction of many interferon-responsive genes. Consistent with this expectation, the true response showed down-regulation of canonical type I interferon-stimulated genes including IFIT2, XAF1, GBP4, GBP5, UBA7, TNFSF10, and CCL5, as well as other interferon- or immune-response-associated genes such as LAMP3, CTSS, and CD74. TranScouter recovered the correct direction for all 25 plotted DEGs. By contrast, scGen reversed the direction of XAF1, GBP4, and CD74, while trVAE reversed GBP4, GBP5, and CD74. These errors are biologically meaningful because they invert the direction of genes whose reduced expression is consistent with attenuated type I interferon signaling after IFNAR2 knockdown. The averaging baseline also made several wrong-direction predictions, although these did not include the same set of canonical interferon-associated genes highlighted above.

In the unseen-perturbation setting, we examined IFNGR2 perturbation in HT-29 cells under IFN-$\gamma$ stimulation (Supplementary Figure 5). IFNGR2 is part of the type II interferon receptor complex, and IFN-$\gamma$ signaling induces transcriptional programs involved in antigen presentation, inflammatory chemokine production, and interferon effector responses. IFNGR2 was withheld from training in this split, so this example tests whether TranScouter can extrapolate to an unseen receptor perturbation in a matching cytokine context. The ground-truth response showed down-regulation of multiple IFN-$\gamma$-responsive genes, including IRF1, NLRC5, TAP1, HLA-B, CXCL11, GBP1, GBP2, GBP4, APOL6, ETV7, and EPSTI1. TranScouter correctly recovered the direction of all 25 plotted DEGs. The correlation-based reference and averaging baseline captured the negative direction of many interferon-response genes, but each reversed several up-regulated genes in the true response, including ANO1 and PADI1. Because HT-29 is an epithelial colorectal cancer cell line, we interpret these up-regulated genes cautiously as cell-line- or cell-state-associated components of the measured response, rather than as direct IFNGR2 targets. This example shows that TranScouter preserved both the attenuation of canonical IFN-$\gamma$ response genes and the reciprocal direction of other DEG-level changes in an unseen-perturbation case.

### Supplementary Note 4: McFaline transferability diagnostic

We also applied the delta-average and sign-match diagnostics to McFaline to test whether the transferable response-change shortcut observed in Jiang24 was also present in this secondary dataset. McFaline is described in Supplementary Note 6. Unlike Jiang24, McFaline did not show a strong transferable directional component. Across 5072 McFaline target pairs, the median sign-match of the averaged response change was 0.50, with an interquartile range of 0.45 to 0.60 (Supplementary Figure 9a). The median target-pair-level mean train-target response correlation was 0.01 (Supplementary Figure 9b). Thus, for a typical McFaline target pair, the average response change transferred from other conditions was close to random directional agreement, and the continuous response profiles were only weakly correlated.

Consistent with this weak transferability, delta-average performed poorly relative to TranScouter on McFaline. In the covariate-balanced overall summaries, TranScouter achieved lower DMR and MSE in both evaluation scenarios (Supplementary Figure 9c): in the seen-perturbation scenario, DMR was 0.35 for TranScouter and 0.50 for delta-average, while MSE was 0.43 for TranScouter and 0.47 for delta-average; in the unseen-perturbation scenario, DMR was 0.36 for TranScouter and 0.51 for delta-average, while MSE was 0.44 for TranScouter and 0.47 for delta-average. At the condition-pair level, TranScouter had lower DMR for 69.8% of McFaline target pairs and lower MSE for 59.6% of target pairs.

The sign-match stratification further supports this interpretation. Low-transferability target pairs dominated McFaline: 3637/5072 target pairs (71.7%) had sign-match $\leq 0.55$, and 5022/5072 target pairs (99.0%) had sign-match $\leq 0.75$. In the sign-match $\leq 0.55$ subset, TranScouter again outperformed delta-average, with median DMR 0.35 versus 0.55 and median MSE 0.32 versus 0.34. High-transferability cases were rare: only 50/5072 target pairs (1.0%) had sign-match $> 0.75$. Therefore, McFaline provides a complementary check on the Jiang24 result: when the transferable response-change shortcut is largely absent, delta-average does not remain competitive, and TranScouter performs more reliably.

### Supplementary Note 5: Rationale for using Jiang24 as the primary benchmark

The main goal of this study is to evaluate cross-condition prediction, where a model is asked to predict perturbation responses in biological conditions that were not observed during training. For this purpose, the relevant unit of dataset diversity is not simply the number of independent dataset files, but the number of biological conditions that can be compared within a coherent experimental framework. A dataset containing many perturbations in a single cell line or basal state is valuable for perturbation extrapolation, but it cannot by itself support evaluation of condition-level generalization.

Jiang24 was selected as the primary benchmark because it provides a relatively broad and internally consistent condition space for genetic perturbation responses. The dataset profiles six cancer cell lines under five cytokine or stimulation conditions, yielding thirty biological conditions measured under a shared experimental design. This structure allows held-out conditions to differ along both cell-line and stimulation axes while reducing the degree to which technical differences between studies are confounded with biological context. In this sense, Jiang24 is more informative for the present task than the phrase "one dataset" might suggest: it contains a collection of internally harmonized condition settings rather than a single biological context.

An alternative strategy would be to combine perturb-seq datasets generated by different studies. Although this could increase the apparent number of conditions, many available genetic perturbation screens are concentrated in a limited set of commonly used immortalized cell lines, so dataset count does not necessarily translate into broad condition coverage. Multi-study integration also introduces several complications for cross-condition evaluation. Perturbed-gene overlap across studies is often incomplete, so restricting analysis to shared targets can discard many informative perturbations. In addition, study-specific batch effects can be substantial, reflecting differences in experimental design, guide libraries, sequencing depth, cell handling, and preprocessing. In a cross-condition benchmark, such effects are especially problematic because dataset identity may become confounded with the biological condition being evaluated. Apparent differences in perturbation response could therefore reflect technical variation rather than true biological context dependence.

We also evaluated TranScouter on McFaline as a secondary multi-condition dataset. McFaline spans fewer biological conditions and is substantially sparser than Jiang24, making it less suitable as the main benchmark but still useful as an additional test of the framework. As larger harmonized perturbation atlases spanning more cell types, stimulation states, and perturbation targets become available, they will enable broader and more definitive evaluation of cross-condition perturbation prediction.

### Supplementary Note 6: Additional benchmark evaluation on McFaline

We used McFaline [1] as a secondary benchmark to examine whether the cross-condition prediction framework could be applied beyond Jiang24. McFaline is a sci-Plex-GxE chemical-genetic CRISPRi screen in glioblastoma cells. For this study, we retained the kinome-wide screen spanning three glioblastoma cell lines (A172, T98G, and U87MG) and five treatment settings: vehicle/no drug, lapatinib targeting EGFR/ERBB signaling, nintedanib targeting PDGFR/FGFR/VEGFR-family receptor tyrosine kinases, trametinib targeting MEK, and zstk474 targeting PI3K. This yielded 15 biological conditions defined by cell line and treatment. The processed benchmark contains 519 non-control perturbation targets. The original data were obtained from GEO under accession GSE225775.

We first visualized the condition structure of the processed McFaline benchmark using a UMAP of unperturbed control cells colored by biological condition (Supplementary Figure 6). The embedding shows that the 15 cell-line-treatment conditions are not transcriptionally identical. The largest separation is associated with cell-line identity, while treatment effects appear as additional shifts within each cell-line background. Thus, McFaline contains meaningful condition variation, but its condition space is less evenly structured than Jiang24 and is dominated more strongly by cell-line differences. McFaline therefore provides a useful secondary test case, but is less suitable as the primary benchmark for the main cross-condition analysis.

McFaline required dataset-specific curation before evaluation. We first followed the PerturBench quality-control workflow [2] to download the original processed objects from GEO, combine the experimental components, standardize metadata, and construct a curated cell-level AnnData object [3]. Starting from this curated cell-level dataset, we retained the main kinome-wide glioblastoma screen and excluded the proof-of-principle A172 MMR/HPRT1 experiments, which used a different perturbation and treatment design. We also excluded cells without assigned guide identities. The original screen included vehicle control, 1 $\mu$M drug exposure, and 10 $\mu$M drug exposure. Because biological conditions in our benchmark were defined by cell line and treatment, including both 1 $\mu$M and 10 $\mu$M exposure under the same treatment label would mix distinct exposure levels within one condition. We therefore retained vehicle control and 10 $\mu$M drug exposure and excluded the 1 $\mu$M exposure.

Unlike Jiang24, McFaline was retained at single-cell resolution after filtering. The retained kinome-wide screen did not provide a stable replicate-level aggregation structure. Finer experimental identifiers produced very small aggregation strata: when cells were grouped by perturbation-condition and experimental identifier, the median stratum contained two cells and more than 99% of strata contained fewer than ten cells. Aggregating only by perturbation-condition would instead collapse each pair into a single expression profile. We therefore used filtered single-cell profiles for this secondary benchmark. Cells were filtered using library-size, detected-gene, and mitochondrial-content criteria, and perturbation-condition groups with too few cells were removed. After preprocessing, each retained perturbation-condition pair contained a median of 26 cells and a mean of 31.3 cells. Differentially expressed genes were then identified within each biological condition for evaluation.

For the McFaline split strategy, we used five condition-level folds. In each split, three biological conditions were held out for testing, two were used for validation, and the remaining ten were used for training. As in the Jiang24 benchmark, we further masked a subset of perturbation targets within each split to evaluate unseen-perturbation prediction. Specifically, 103 perturbation targets were withheld from training in each split and used to define the unseen-perturbation evaluation scenario.

McFaline results are summarized in Supplementary Figure 7 and Supplementary Figure 8. In the seen-perturbation setting, TranScouter remained competitive with scGen but did not outperform it on MSE. Averaged across conditions, the median MSE on DEGs was 0.32 for TranScouter, 0.31 for scGen, and 0.57 for the averaging baseline. The corresponding median DMR values were 0.35 for TranScouter, 0.35 for scGen, and 0.45 for the averaging baseline. In the unseen-perturbation setting, TranScouter achieved the lowest overall error among the evaluated methods, with a median MSE of 0.32 compared with 0.32 for the correlation-based reference and 0.57 for the averaging baseline. Although the MSE difference between TranScouter and the correlation-based reference was small, TranScouter showed a clearer advantage on directionality, with a median DMR of 0.35 compared with 0.50 for the correlation-based reference and 0.45 for the averaging baseline. These results support the main conclusion from Jiang24 that perturbation representations can provide useful signal for targets not observed during training, while suggesting that performance may be more modest when substantially fewer training conditions are available; in McFaline, each split used 10 training conditions, compared with 25 training conditions in the Jiang24 benchmark. trVAE was not included in the McFaline results because training did not complete within 15 days. CellOracle was also not evaluated on McFaline because only two perturbation targets, TAF1 and TRIM28, were transcription factors represented in the CellOracle GRN-based perturbation workflow, leaving too few applicable perturbations for a meaningful comparison.

We further examined representative perturbation-condition pairs to assess whether the directional advantages corresponded to interpretable transcriptional responses. In the seen-perturbation setting, we examined ADCK3 perturbation in A172 cells treated with lapatinib (Supplementary Figure 10). ADCK3, also known as COQ8A, is linked to mitochondrial coenzyme Q biology, while lapatinib inhibits EGFR/ERBB signaling and can alter growth-associated signaling and downstream metabolic programs. The true response included metabolic genes including PKM, lipid-associated genes including SCD and FADS2, and cytoskeleton-associated genes such as CDC42EP3 and DBN1. We interpret these genes as response markers in this treatment context rather than as evidence for a direct pathway from ADCK3 to each DEG. TranScouter recovered the correct direction for all 20 plotted DEGs, whereas scGen reversed several genes including SCD, FADS2, PKM, CDC42EP3, and DBN1, and the averaging baseline also reversed SCD and FADS2. This example illustrates that, even when scGen performs similarly on aggregate McFaline metrics, TranScouter can better preserve the direction of specific DEG-level responses in individual cases.

In the unseen-perturbation setting, we examined MAP4K4 perturbation in U87MG cells treated with nintedanib (Supplementary Figure 11). Nintedanib inhibits multiple receptor tyrosine kinases, including VEGFR, FGFR, and PDGFR family members, while MAP4K4 is a kinase implicated in stress-activated MAPK signaling, cytoskeletal regulation, and cell-migration-related programs. In this case, TranScouter recovered the correct direction for 19 of the 20 plotted DEGs, missing only SAMD4A. Among the genes where the reference methods predicted the wrong direction, several have functional annotations that may be relevant to cellular-state changes in this perturbation-treatment context, including CALR, which is associated with endoplasmic-reticulum protein folding, CD63, which is linked to endolysosomal compartments, and NUMB, WNK1, and SH2B3, which participate in signaling or adaptor functions. The correlation-based reference reversed many of these genes, and the averaging baseline also made multiple wrong-direction predictions. These examples should be viewed as qualitative illustrations rather than mechanistic validation, but they show cases in which TranScouter better preserved the direction of biologically interpretable DEG-level responses.

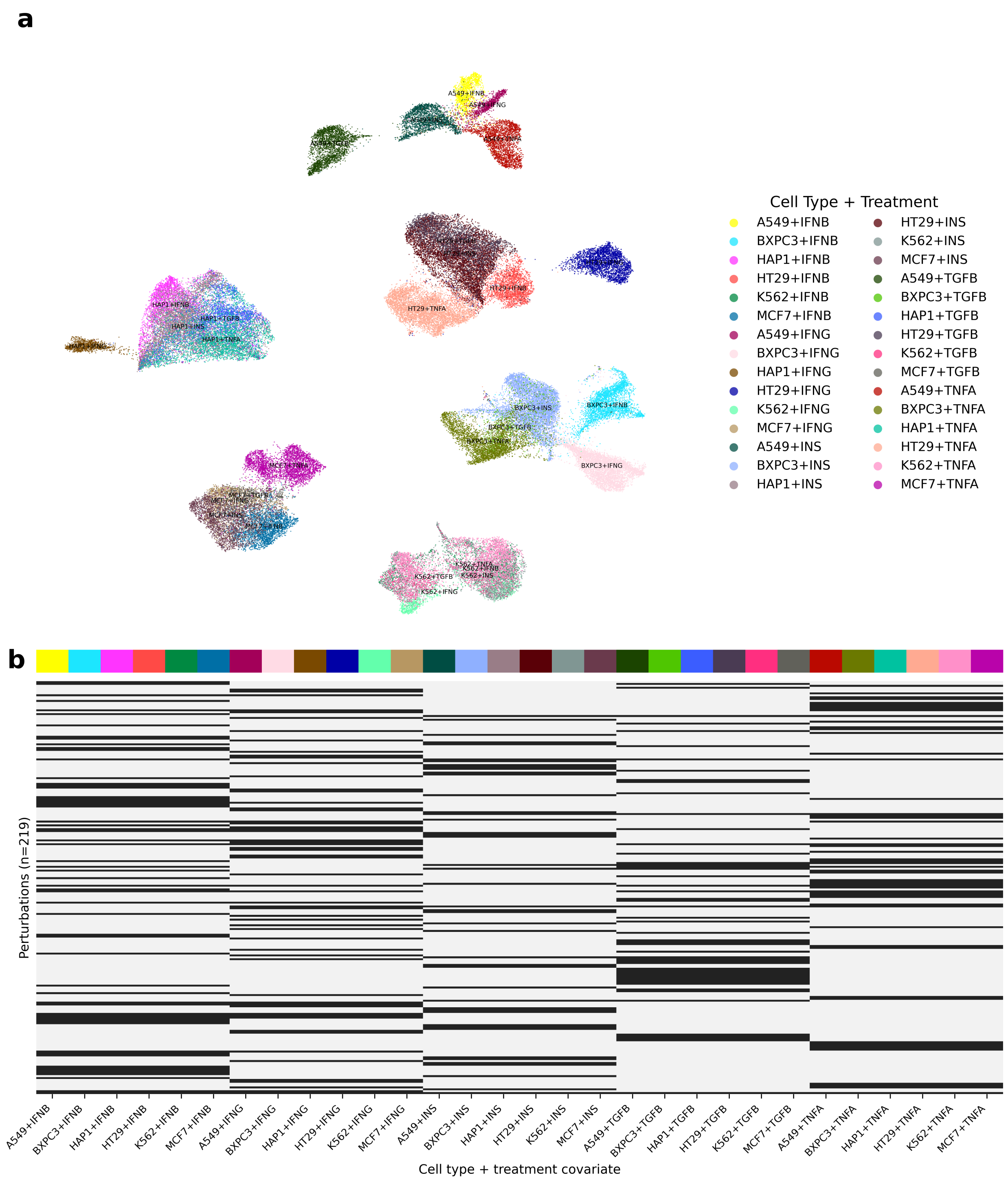


Figure 1: **Biological condition structure and perturbation coverage in Jiang24.** **a**, UMAP visualization of 84,269 unaggregated control cells, colored by biological condition defined as cell line plus treatment. The separation among many cell line-treatment groups indicates that the benchmark contains transcriptionally distinct baseline conditions. **b**, Perturbation-by-condition availability heatmap for the processed Jiang24 benchmark. Columns correspond to the 30 biological conditions and rows correspond to perturbation labels, including the control label. Black marks indicate observed perturbation-condition combinations.

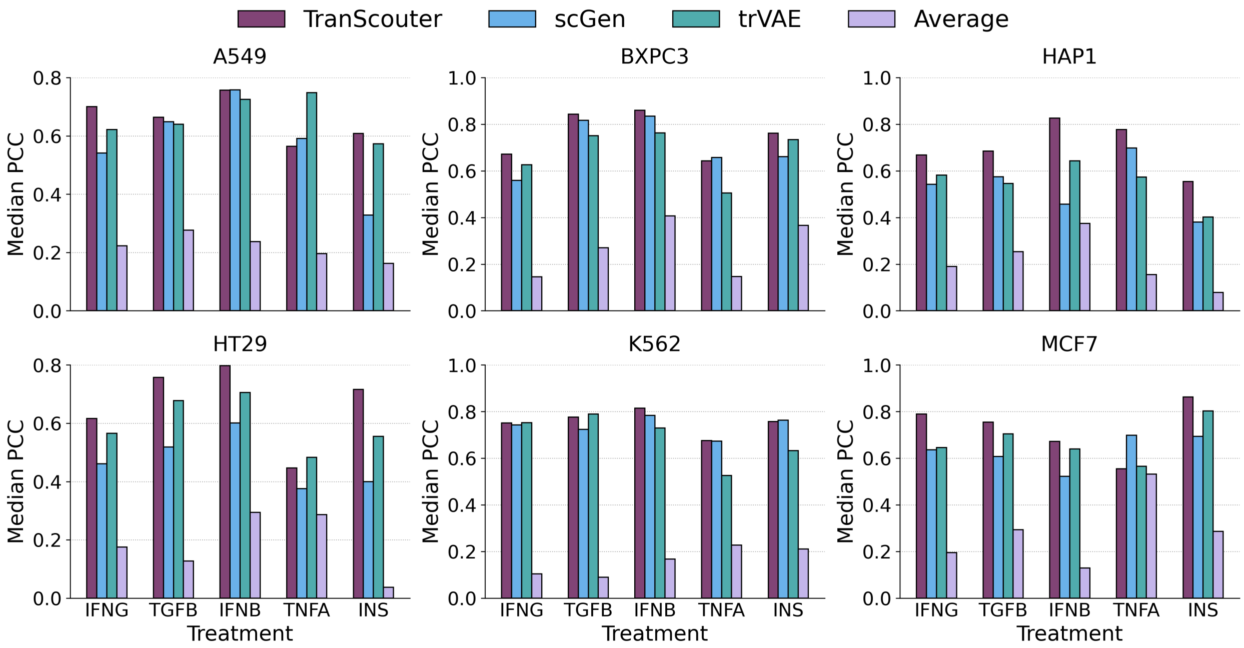


Figure 2: **PCC analysis in the Jiang24 seen-perturbation setting.** Median Pearson correlation coefficient (PCC) on DEGs for held-out biological conditions. PCC was computed for perturbation-condition pairs with more than ten DEGs and summarized by condition. TranScouter is compared with scGen, trVAE, and the perturbation-averaging baseline.

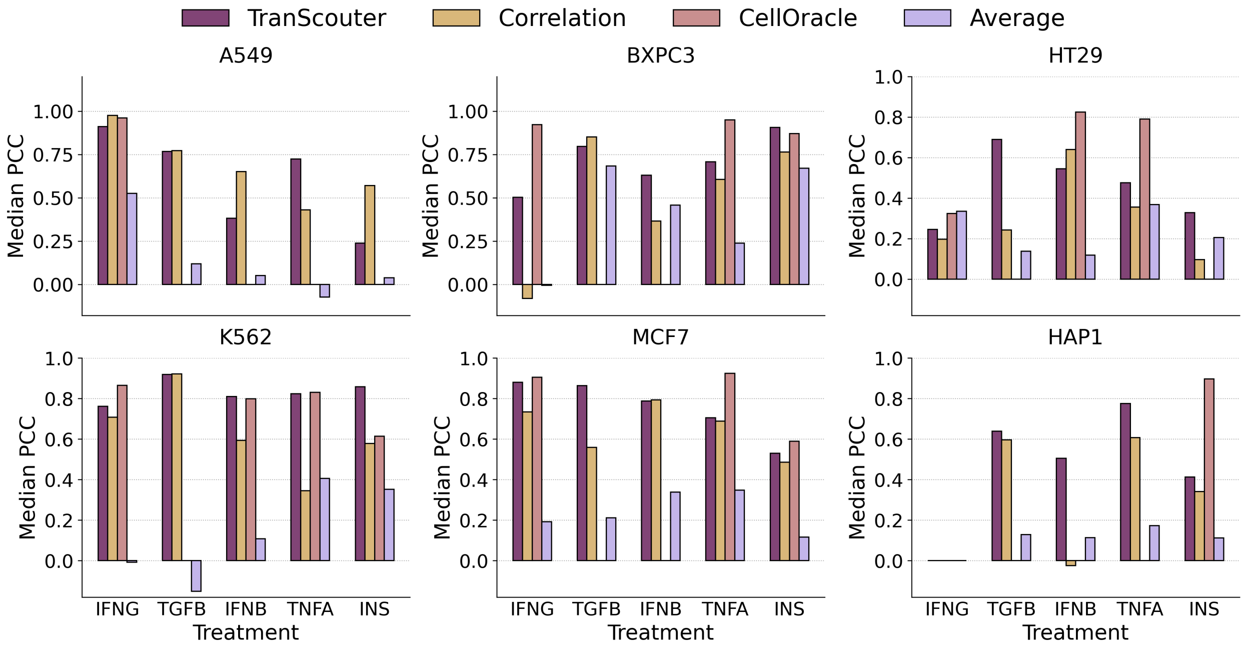


Figure 3: **PCC analysis in the Jiang24 unseen-perturbation setting.** Median Pearson correlation coefficient (PCC) on DEGs for held-out biological conditions and perturbation targets withheld from training. PCC was computed for perturbation-condition pairs with more than ten DEGs and summarized by condition. Missing CellOracle bars indicate conditions where CellOracle could not be evaluated because the relevant perturbations were outside the TF scope of its GRN-based simulation framework.

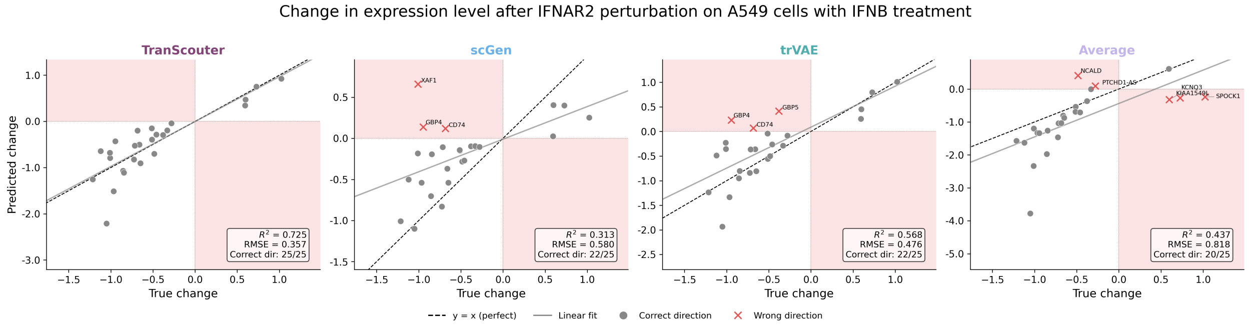


Figure 4: **Example seen-perturbation prediction for IFNAR2 perturbation under IFN-$\beta$ stimulation.** Predicted versus observed DEG-level expression changes for IFNAR2 perturbation in A549 cells treated with IFN-$\beta$. Red shaded quadrants indicate predictions with the opposite sign from the true expression change. Panel annotations report squared Pearson correlation $R^2$, root mean square error (RMSE), and the number of DEGs with correctly predicted direction.

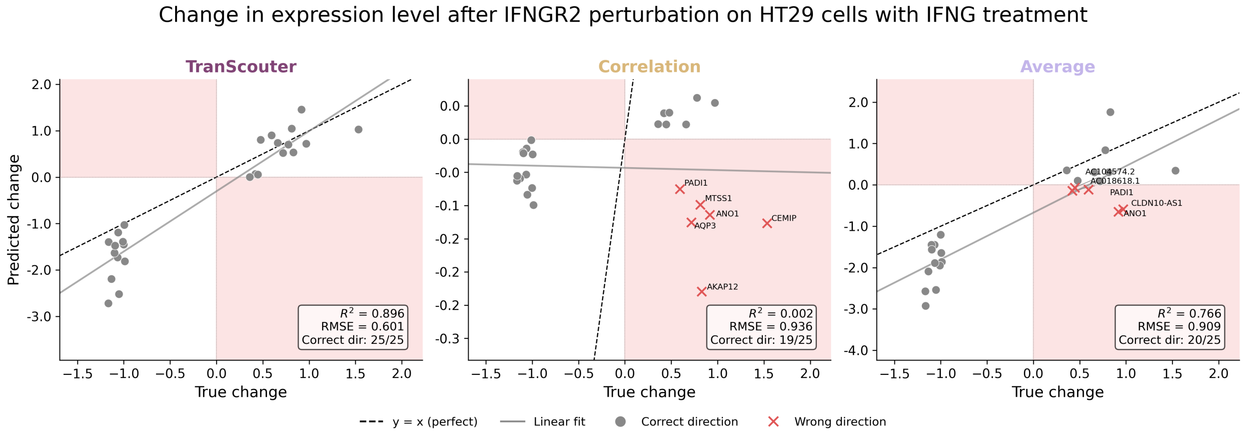


Figure 5: **Example unseen-perturbation prediction for IFNGR2 perturbation under IFN-$\gamma$ stimulation.** Predicted versus observed DEG-level expression changes for IFNGR2 perturbation in HT-29 cells treated with IFN-$\gamma$. IFNGR2 was withheld from training in this split. Red shaded quadrants indicate predictions with the opposite sign from the true expression change. Panel annotations report squared Pearson correlation $R^2$, root mean square error (RMSE), and the number of DEGs with correctly predicted direction.

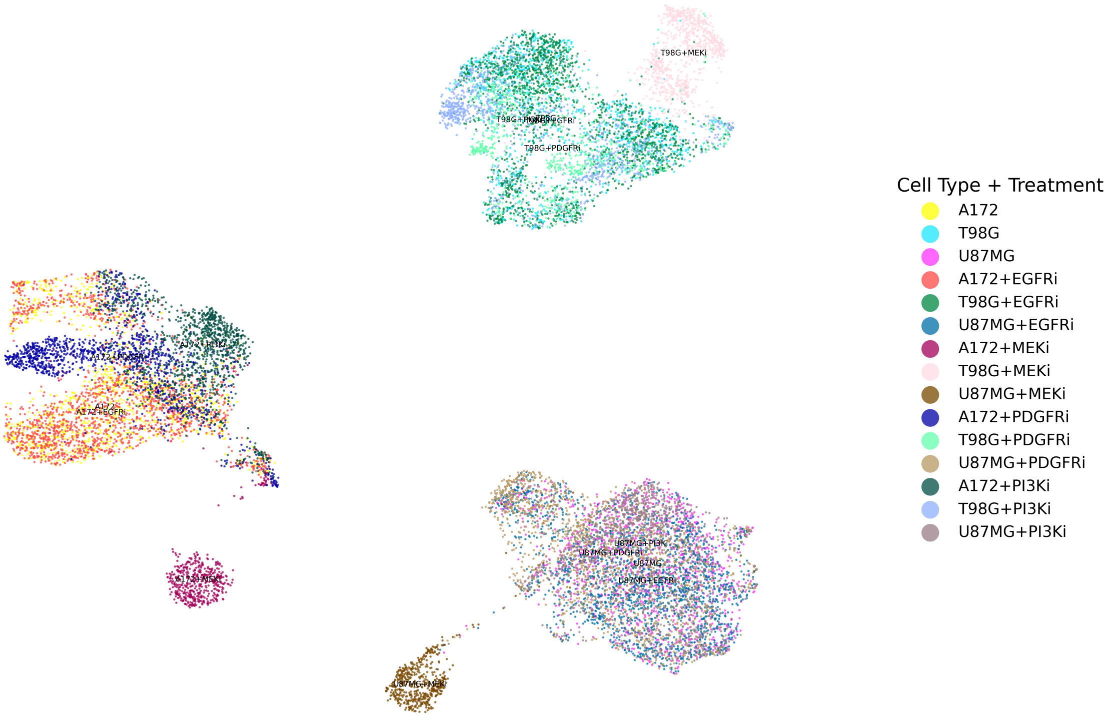


Figure 6: **Biological condition structure in the McFaline secondary benchmark.** UMAP visualization of unperturbed control cells in the processed McFaline benchmark, colored by biological condition defined as cell line plus treatment. Treatment labels are shown using inhibitor-class abbreviations. The embedding shows that McFaline contains transcriptional variation across cell-line-treatment conditions, with separation driven primarily by cell-line identity and additional treatment-associated shifts within each cell-line background.

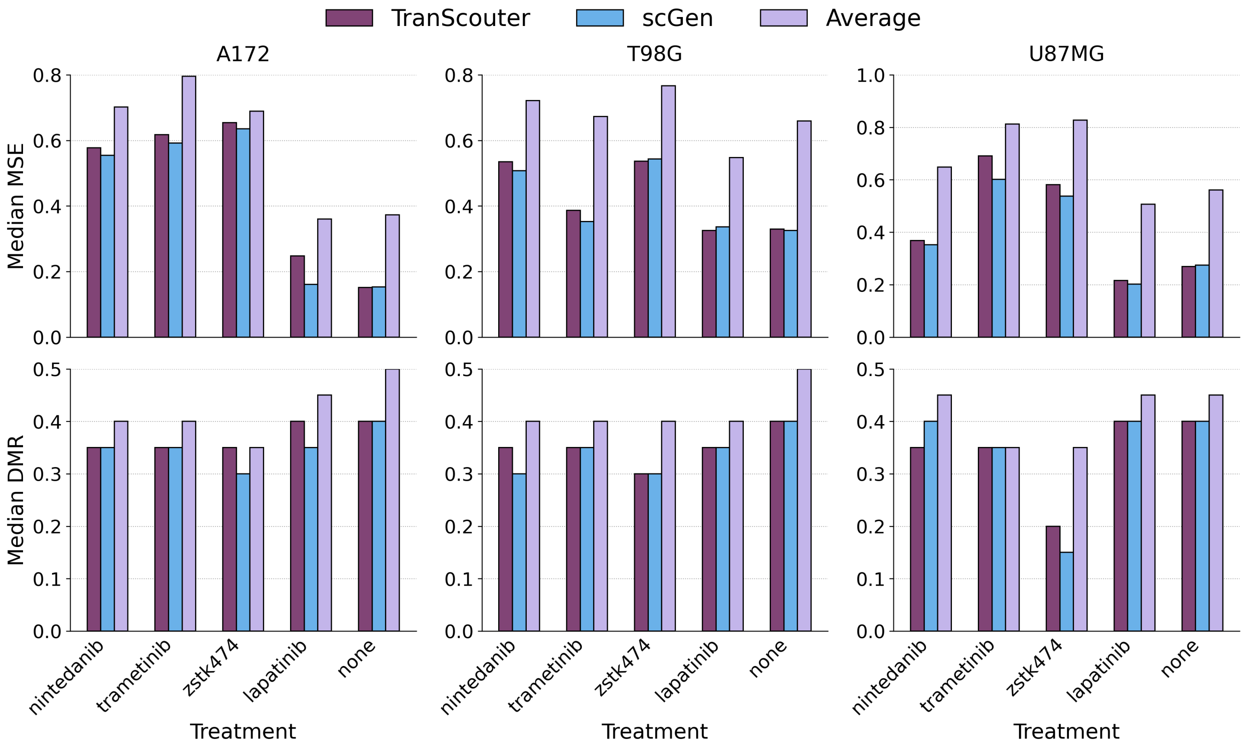


Figure 7: **McFaline performance in the seen-perturbation setting.** Median MSE and DMR on DEGs for held-out biological conditions in the McFaline secondary benchmark. Results are grouped by cell line and treatment. TranScouter is compared with scGen and the perturbation-averaging baseline.

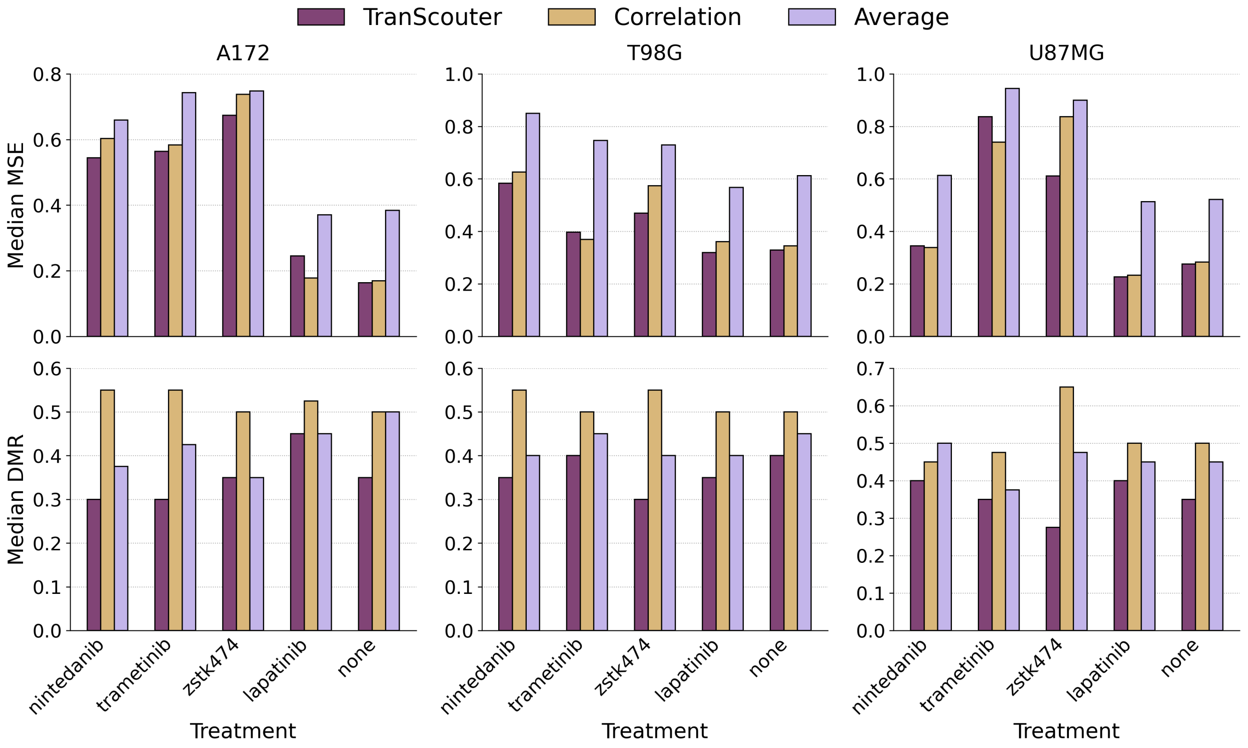


Figure 8: **McFaline performance in the unseen-perturbation setting.** Median MSE and DMR on DEGs for held-out biological conditions and perturbation targets withheld from training in the McFaline secondary benchmark. TranScouter is compared with the correlation-based reference and the perturbation-averaging baseline.

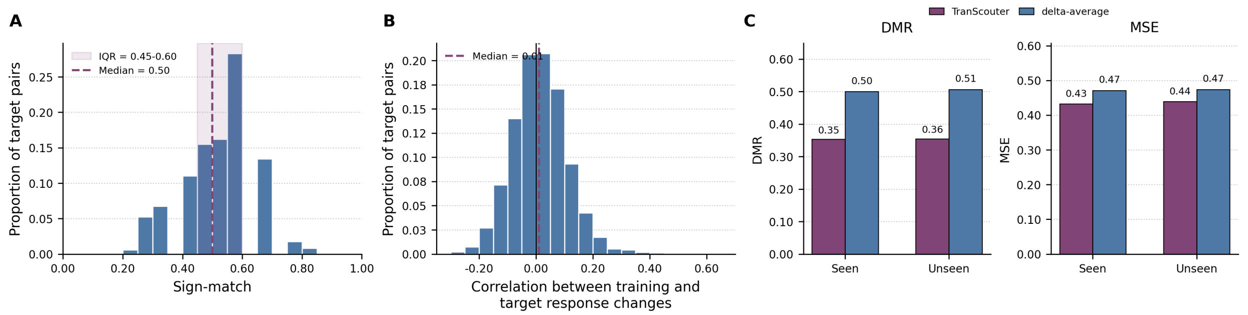


Figure 9: **McFaline transferability diagnostics.** **a**, Distribution of sign-match scores across McFaline target perturbation-condition pairs. The shaded region marks the interquartile range and the dashed line marks the median. **b**, Distribution of target-pair-level mean train-target response correlations. For each target pair, the response correlation is first computed between each available training-condition response change and the target-condition response change on the DEG set, and then averaged across training conditions. The dashed line marks the median. **c**, Covariate-balanced overall DMR and MSE for TranScouter and delta-average in the seen- and unseen-perturbation scenarios. Lower values indicate better performance.

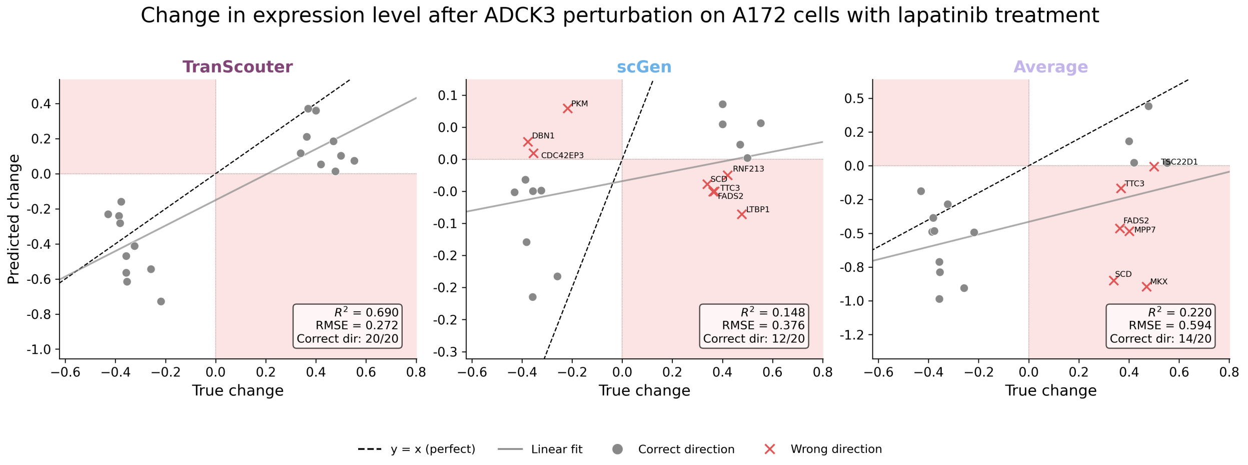


Figure 10: **Example seen-perturbation prediction in McFaline.** Predicted versus observed DEG-level expression changes for ADCK3 perturbation in A172 cells treated with lapatinib. Red shaded quadrants indicate predictions with the opposite sign from the true expression change. Panel annotations report squared Pearson correlation $R^2$, root mean square error (RMSE), and the number of DEGs with correctly predicted direction.

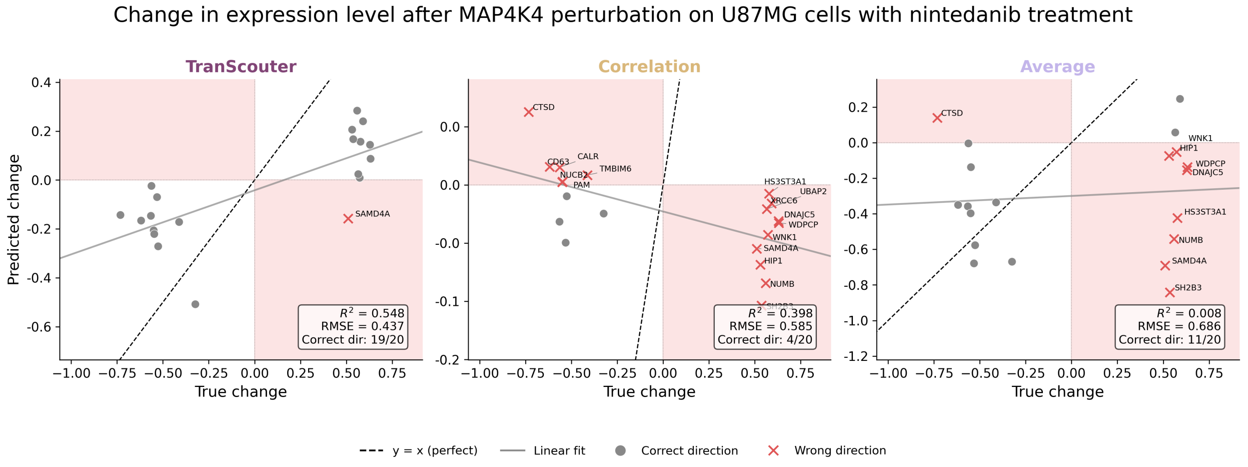


Figure 11: **Example unseen-perturbation prediction in McFaline.** Predicted versus observed DEG-level expression changes for MAP4K4 perturbation in U87MG cells treated with nintedanib. MAP4K4 was withheld from training in this split. Red shaded quadrants indicate predictions with the opposite sign from the true expression change. Panel annotations report squared Pearson correlation $R^2$, root mean square error (RMSE), and the number of DEGs with correctly predicted direction.
